# The genomics of mimicry: gene expression throughout development provides insights into convergent and divergent phenotypes in a Müllerian mimicry system

**DOI:** 10.1101/706671

**Authors:** Adam M M Stuckert, Mathieu Chouteau, Melanie McClure, Troy M LaPolice, Tyler Linderoth, Rasmus Nielsen, Kyle Summers, Matthew D MacManes

## Abstract

A common goal in evolutionary biology is to discern the mechanisms that produce the astounding diversity of morphologies seen across the tree of life. Aposematic species, those with a conspicuous phenotype coupled with some form of defense, are excellent models to understand the link between vivid color pattern variations, the natural selection shaping it, and the underlying genetic mechanisms underpinning this variation. Mimicry systems in which multiple species share the same conspicuous phenotype can provide an even better model for understanding the mechanisms of color production in aposematic species, especially if comimics have divergent evolutionary histories. Here we investigate the genetic mechanisms by which vivid color and pattern are produced in a Müllerian mimicry complex of poison frogs. We did this by first assembling a high-quality de novo genome assembly for the mimic poison frog *Ranitomeya imitator*. This assembled genome is 6.8 Gbp in size, with a contig N50 of 300 Kbp and 93% of expected tetrapod genes. We then leveraged this genome to conduct gene expression analyses throughout development of four color morphs of *R. imitator* and two color morphs from both *R. fantastica* and *R. variabilis* which *R. imitator* mimics. We identified a large number of pigmentation and patterning genes that are differentially expressed throughout development, many of them related to melanocyte development, melanin synthesis, iridophore development, and guanine synthesis. In addition, we identify the pteridine synthesis pathway (including genes such as *qdpr* and *xdh*) as a key driver of the variation in color between morphs of these species. Finally, we hypothesize that genes in the keratin family are important for producing different structural colors within these frogs.

## Introduction

The diversity of animal coloration in the natural world has long been a focus of investigation in evolutionary biology (Beddard, 1892; Fox, 1936; Gray & McKinnon, 2007; Longley, 1917). Color phenotypes can be profoundly shaped by natural selection, sexual selection, or both. Further, these color phenotypes are often under selection from multiple biotic (e.g. competition, predation) and abiotic (e.g. temperature, salinity) factors (Rudh & Qvarnström, 2013). The mechanisms underlying color and pattern phenotypes are of general interest because they can help explain the occurrence of specific evolutionary patterns, particularly in systems where these phenotypes embody key adaptations driving biological diversification.

Adaptive radiations in aposematic species (those species which couple conspicuous phenotypes with a defense), provide an example of such phenotypes under strong selection (Kang et al., 2017; Ruxton, Sherratt, & Speed, 2004; Sherratt, 2006, 2008). In these biological systems, geographically heterogeneous predation produces rapid diversification of color and pattern within a species or group of species. This produces a diversity of polytypic phenotypes (defined as distinct defensive warning colorations in distinct localities) that are maintained geographically, with each population characterized by a unique phenotype that deters predators. This spatial mosaic of local adaptations maintained by the strong stabilizing selection exerted by predators, also results in convergence of local warning signals in unrelated species. Examples of impressive diversification within species and mimetic convergence between species have been documented in many biological systems, including *Heliconius* butterflies (Mallet & Barton, 1989), velvet ants (Wilson et al., 2015), millipedes (Marek & Bond, 2009), and poison frogs (Stuckert, Saporito, Venegas, & Summers, 2014; Stuckert, Venegas, & Summers, 2014; Symula, Schulte, & Summers, 2001; Twomey et al., 2013), to name only a few of the documented examples of diverse aposematic phenotypes (Briolat et al., 2019).

Understanding the genomic architecture of adaptive phenotypic diversification that underpins these diversification events has therefore been of substantial importance to the field of evolutionary biology (Hodges & Derieg, 2009). This knowledge is crucial to examine selection on color pattern phenotypes and determine the mechanisms by which convergence occurs at the molecular level. The majority of our knowledge of the genomics of warning signals and mimicry comes from butterflies of the genus *Heliconius*. In these organisms, a handful of key genetic loci of large phenotypic effect controls the totality of phenotypic variation observed within populations of the same species, but also explains the mimicry convergence between distinct species (e.g., *WntA* (Martin et al., 2012) and *optix* (Reed et al., 2011; Supple et al., 2013)), though there are many others likely involved as well (Kronforst & Papa, 2015). While *Heliconius* butterflies are excellent subjects for the study of mimicry, characterizing the genetics of mimicry in a phylogenetically distant system is critically important to determining whether adaptive radiations in warning signals and mimitic relationships are generally driven by a handful of key loci.

Here we investigate the genetics of convergent color phenotypes in *Ranitomeya* from Northern Peru. In this system, the mimic poison frog *(Ranitomeya imitator,* Dendrobatidae; (Schulte, 1986) underwent a rapid adaptive radiation, such that it mimics established congenerics (*R. fantastica, R. summersi*, and *R. variabilis*), thereby “sharing” the cost of educating predators—a classic example of Müllerian mimicry (Stuckert, Saporito, et al., 2014; Stuckert, Venegas, et al., 2014; Symula et al., 2001; Symula, Schulte, & Summers, 2003). This is a powerful system for evolutionary study of color patterns, as the different *R. imitator* color morphs have undergone adaptive divergence to converge on shared phenotypes with the species that *R. imitator* mimics.

Furthermore, despite new genomic data shedding critical insights into the mechanisms of color production in amphibians in general (Burgon et al., 2020), and poison frogs in particular (Rodríguez, Mundy, Ibáñez, & Pröhl, 2020; Stuckert et al., 2019; Twomey, Johnson, Castroviejo-Fisher, & Van Bocxlaer, 2020), our knowledge of amphibian genomics is critically behind that of other taxa, both in terms of genomic resources as well as our ability to make inferences from these resources. This is largely due to the challenging nature of these genomes, most of which are extremely large and rich with repeat elements that have proliferated throughout the genome. This makes many amphibians a nearly intractable system for in-depth genomic analyses (Funk, Zamudio, & Crawford, 2018; Rogers et al., 2018). As a result, there is a dearth of publically available amphibian genomes (14 anuran species as of September 1st, 2020). In fact, many of the available genomes are from a single group of frogs with a genome size of less than 1 Gbp, which is on the lower bound of known amphibian genome size (Funk et al., 2018).

We investigated the genetics of Müllerian mimicry by first generating a high quality 6.8 Gbp *de novo* genome for the mimic poison frog, *Ranitomeya imitator*. This is an important new resource for amphibian biologists, as it fills a substantial gap in the amphibian phylogeny and enables for more detailed comparative work. We then utilized this high quality *Ranitomeya imitator* genome to examine gene expression patterns using RNA sequencing from early tadpole development all the way through to the end of metamorphosis in both the mimic (*R. imitator*) and model species (*R. fantastica* and *R. variabilis*). As such, we were able to keep track of patterns of expression in genes responsible for color throughout development both between color morphs, and between species. Color patterns within these species begin to appear early during development when individuals are still tadpoles, which is consistent with observations that chromatophores develop from the neural crest early during embryonic development (DuShane, 1935). This comparative genomic approach allowed us to carefully examine the genes and gene networks responsible for diversification of color patterns and mimicry in poison frogs.

## Methods

### IACUC statements

#### R. imitator

Animal use and research comply with East Carolina University’s IACUC (AUP #D281) and the University of New Hampshire’s IACUC (AUP #180705).

#### *R. fantastica* and *R. variabilis*

The protocol for *R. fantastica* and *R. variabilis* sample collection was approved by the Peruvian Servicio Forestal y de Fauna Silvestre through the authorization number 232-2016-SERFOR/DGGSPFFS and export permit N° 17PE 001718 and the authorization from the French Direction de l’Environnement, de l’Agriculture, de l’Alimentation et de la forêt en Guyane number 973-ND0073/SP2000116-13.

### Data accessibility

All read data, *de novo* transcriptome assemblies, our *de novo* genome assembly, and our annotations are archived with the European Nucleotide Archive (accession number PRJEB28312; https://www.ebi.ac.uk/ena/browser/view/PRJEB28312). Code for assemblies, annotation, and all subsequent analyses are all available on GitHub (https://github.com/AdamStuckert/Ranitomeya_imitator_genome).

### Generating a *de novo* genome for *Ranitomeya imitator*

#### Genome Sequencing Approach

##### 10X Chromium

A single, likely male, *R. imitator* of the ‘intermedius’ morph from the pet trade was euthanized and high molecular weight DNA was extracted from liver tissue using the QIAGEN Blood & Cell Culture DNA Kit. 10X Genomics Chromium Genome library (Weisenfeld, Kumar, Shah, Church, & Jaffe, 2017) was prepared by the DNA Technologies and Expression Analysis Cores at the University of California Davis Genome Center and sequenced on an Illumina HiSeq X by Novogene Corporation (Mudd et al. in prep).

##### Long read (Nanopore and PacBio) sequencing

Captive bred frogs from the pet trade that originated from the Tarapoto region (green-spotted morph) were euthanized and the skin and gastrointestinal tract was removed in order to reduce potential contamination from skin and gut microbial communities. Each frog was dissected into 8 approximately equal chunks of tissue and DNA was extracted using a Qiagen Genomic Tip extraction kit. DNA concentration was quantified with a Qubit 3.0 and fragment length was assessed with a TapeStation using a D1000 kit.

For Nanopore sequencing we prepared libraries for direct sequencing via Oxford Nanopore using a LSK-109 kit. Samples were loaded onto either R9 or R10 flowcells, which yielded minimum throughput. We basecalled raw fast5 files from Nanopore sequencing using the “read_fast5_basecaller.py” script in the ONT Albacore Sequencing Pipeline Software version 2.3.4.

For Pacific Biosystems (PacBio) sequencing we used a Circulomics short-read eliminator kit to size select extracted DNA. After this, we sent ~15 μg of high molecular weight DNA to the Genomics Core Facility in the Icahn School of Medicine at Mt. Sinai for library preparations and sequencing. Here libraries were prepared with 20-25 kb inserts and were sequenced on three SMRTcell 8M cells on a Pacific Biosciences Sequel II.

##### Genome assembly

We took a multifaceted approach to constructing the *Ranitomeya imitator* genome, which contained iterative scaffolding steps. We detail our general approach here, and have a graphical depiction of this in Figure 2. We converted the “good” subreads in “.bam” format into fasta files using the *samtools* “fasta” command (Heng Li et al., 2009). We then used the contig assembler wtdbg2 version 2.5 *(Ruan & Li, 2019)* to create our initial assembly and create consensus contigs using only the PacBio data. Wtdbg2 uses a fuzzy de bruijn graph approach to combine reads into contigs. After producing this assembly, we polished (error corrected) the assembly using Illumina 10X reads. To do this we first mapped the Illumina short reads to our assembly using BWA version 0.7.17-r1188 (H. Li & Durbin, 2009). Polishing was then conducted using the software Pilon version 1.22 (Walker et al., 2014).

We then used Arcs version 3.82 (Coombe et al., 2018) to scaffold our existing assembly using Illumina 10X data. Prior to doing this, we used the “basic” function within the program Longranger (Marks et al., 2019) to trim, error correct, and barcode the 10X data. We then used the “arks.mk” makefile provided in the Arcs GitHub page (https://github.com/bcgsc/arcs/blob/master/Examples/arcs-make), to run the Arcs software. This makefile aligns the barcoded 10X data from Longranger using BWA, then runs Arcs to scaffold our assembly. After scaffolding our assembly with the 10X data, we ran one round of Cobbler (Warren, 2016) to gap-fill regions of unknown nucleotides (depicted as “N”s) in our assembly. Our input data was the full set of our PacBio and Nanopore data, which was mapped to our assembly using minimap2 version 2.10-r761 (Heng Li, 2018) with no preset for sequencing platform. We then ran RAILS version 5.26.1 (Warren, 2016) to scaffold again, with the same input data as for Cobbler. We aligned the long-read data with Minimap2 because it maps a higher proportion of long reads than other aligners and we used BWA because it is more accurate for short-read data (Heng Li, 2018). After this we polished our assembly again by mapping Illumina reads with BWA and Pilon.

We then attempted to fill in gaps within our assembly using LR Gapcloser (Xu et al., 2019). We ran this program iteratively, first using the Nanopore data and specifying the “-s n” flag. We then filled gaps using LR Gapcloser with PacBio reads and the “-s p” flag. Following this, we again polished our assembly using BWA and Pilon.

We examined genome quality in two main ways. First we examined the presence of genic content in our genome using Benchmarking Universal Single-Copy Orthologs version 3.0.0 (BUSCO; (Simão, Waterhouse, Ioannidis, Kriventseva, & Zdobnov, 2015) using the tetrapod database (tetrapoda_odb9; 2016-02-13). BUSCO version 4.0 was released during this project, so we also examined our final assembly with BUSCO version 4.0.6 and the tetrapod database (tetrapoda_odb10; 2019-11-20). Our second method of genome examination was genome contiguity. We used the assemblathon perl script (https://github.com/KorfLab/Assemblathon) to calculate overall numbers of scaffolds and contigs as well as contiguity metrics such as N50 for both.

##### Repeats and genome annotation

We modeled genomic repeats with Repeat Modeler version 1.0.8 (Smit & Hubley, 2008-2015) using RepBase database 20170127. We used the classified consensus output from Repat Modeler as input to Repeat Masker version 4.1.0 (Smit, Hubley, & Green, 2013-2015) with the *Homo* database specified. We used a combined database of Dfam_3.1 and RepBase-20181026 (Bao, Kojima, & Kohany, 2015).

We annotated our genome using Maker version 3.01.02 (Campbell, Holt, Moore, & Yandell, 2014). We used transcript evidence from *R. imitator* to aid in assembly (“est2genome=1”). For details on transcript evidence used to annotate this genome please see the Appendix. We annotated the genome using both the draft genome as well as a repeat masked genome from Repeat Modeler and Repeat Masker in order to identify the best annotation of our genome.

We used BUSCO version 4.0.6 (Simao *et al.* 2015) on the final version of the *R. imitator* genome to locate the duplicated regions within the genome. BUSCO searches a collated database of benchmarked universal single copy orthologs to measure completeness of the genome. BUSCO identifies whether these single copy orthologs are present in the genome and if they are complete or fragmented. Additionally, it identifies whether these orthologs are present in a single copy or duplicated in the genome. Given the high proportion of duplicated orthologs present in our genome, we proceeded to investigate the read support for each of these duplicated regions across our sequencing technologies. We aligned PacBio data using Minimap2 version 2.10 (Heng Li, 2018). After we aligned the data, we used SAMtools version 1.10 (Heng Li et al., 2009) to generate the depth at each base in the genome for data from each sequencing platform. We wrote a custom Python script (see code availability statement for link to code repository) to extract the depth at the duplicated regions of the genome. Additionally, we extracted the genic regions identified as single copy genes by BUSCO in the same fashion. This allowed us to examine if there were any discrepancies between the duplicates and the rest of the genomic sequence. For each region in both the single copy and duplicated genes we calculated mean and standard deviation. We identified regions that were outliers in sequencing depth by identifying genes that had average sequencing depth that was outside of the average genome-wide sequencing depth plus two standard deviations, or below 10x coverage. We did this because average coverage varied dramatically at a per-base scale (PacBio genome wide coverage: average = 34.5 ± 79.1 sd). We view this as a method of detecting duplicated regions that have plausibly been incorrectly assembled (i.e., spuriously collapsed or duplicated), and is not definitive *per se.*

### IDENTIFICATION OF COLOR PATTERN CANDIDATE GENES

#### Gene expression sample preparation

##### Ranitomeya imitator

The initial breeding stock of *Ranitomeya imitator* was purchased from Understory Enterprises, LLC (Chatham, Canada). Frogs used in this project represent captive-bred individuals sourced from the following wild populations: Baja Huallaga (yellow-striped morph), Sauce (orange-banded), Tarapoto (green-spotted), and Varadero (redheaded; see Figure 1). We raised tadpoles on a diet of Omega One Marine Flakes fish food mixed with Freeze Dried Argent Cyclop-Eeze, which they received three times a week, with full water changes twice a week until sacrificed for analyses at 2, 4, 7, and 8 weeks of age. We sequenced RNA from a minimum of three individuals at each time point from the Sauce, Tarapoto, and Varadero populations (except for Tarapoto at 8 weeks), and two individuals per time point from the Huallaga population. Individuals within the same time points were sampled from different family groups (Table 1).

**Table 1.** Repeat elements classified by Repeat Masker. Most repeats fragmented by insertions or deletions were counted as a single element.

| Element type |  |  | Number of elements | Total length of elements (bp) | Percentage of genomic sequence |
| --- | --- | --- | --- | --- | --- |
| Retroelements |  |  | 1108460 | 1247087328 | 18.36 |
|  | SINEs |  | 5440 | 428468 | 0.01 |
|  | Penelope |  | 60001 | 32685796 | 0.48 |
|  | LINEs |  | 655737 | 671087603 | 9.88 |
|  |  | CRE/SLACS | 0 | 0 | 0 |
|  |  | L2/CR1/Rex | 241795 | 194226476 | 2.86 |
|  |  | R1/LOA/Jockey | 0 | 0 | 0 |
|  |  | R2/R4/NeSL | 1089 | 644862 | 0.01 |
|  |  | RTE/Bov-B | 8249 | 5177385 | 0.08 |
|  |  | L1/CIN4 | 343520 | 436013718 | 6.42 |
|  | LTR elements |  | 447283 | 575571257 | 8.47 |
|  |  | BEL/Pao | 3504 | 2966778 | 0.04 |
|  |  | Gypsy/DIRS1 | 340203 | 505961953 | 7.45 |
|  |  | Retroviral | 85837 | 44644563 | 0.66 |
| DNA transposons |  |  | 703907 | 423320187 | 6.23 |
|  | hobo-Activator |  | 144891 | 93261380 | 1.37 |
|  | Tc1-IS630-Pogo |  | 504231 | 296915689 | 4.37 |
|  | En-Spm |  | 0 | 0 | 0 |
|  | MuDR-IS905 |  | 0 | 0 | 0 |
|  | PiggyBac |  | 10239 | 8260072 | 0.12 |
|  | Tourist/Harbinger |  | 19834 | 14807248 | 0.22 |
|  | Other (Mirage, P-element, Transib) |  | 0 | 0 | 0 |
| Rolling-circles |  |  | 4811 | 2310034 | 0.03 |
| Unclassified |  |  | 5605837 | 1743383014 | 25.67 |
| Total interspersed repeats |  |  |  | 3413790529 | 50.26 |
| Small RNA |  |  | 0 | 0 | 0 |
| Satellites |  |  | 0 | 0 | 0 |
| Simple repeats |  |  | 1930599 | 246155996 | 3.62 |
| Low complexity |  |  | 211720 | 14981007 | 0.22 |

**Figure 1.**
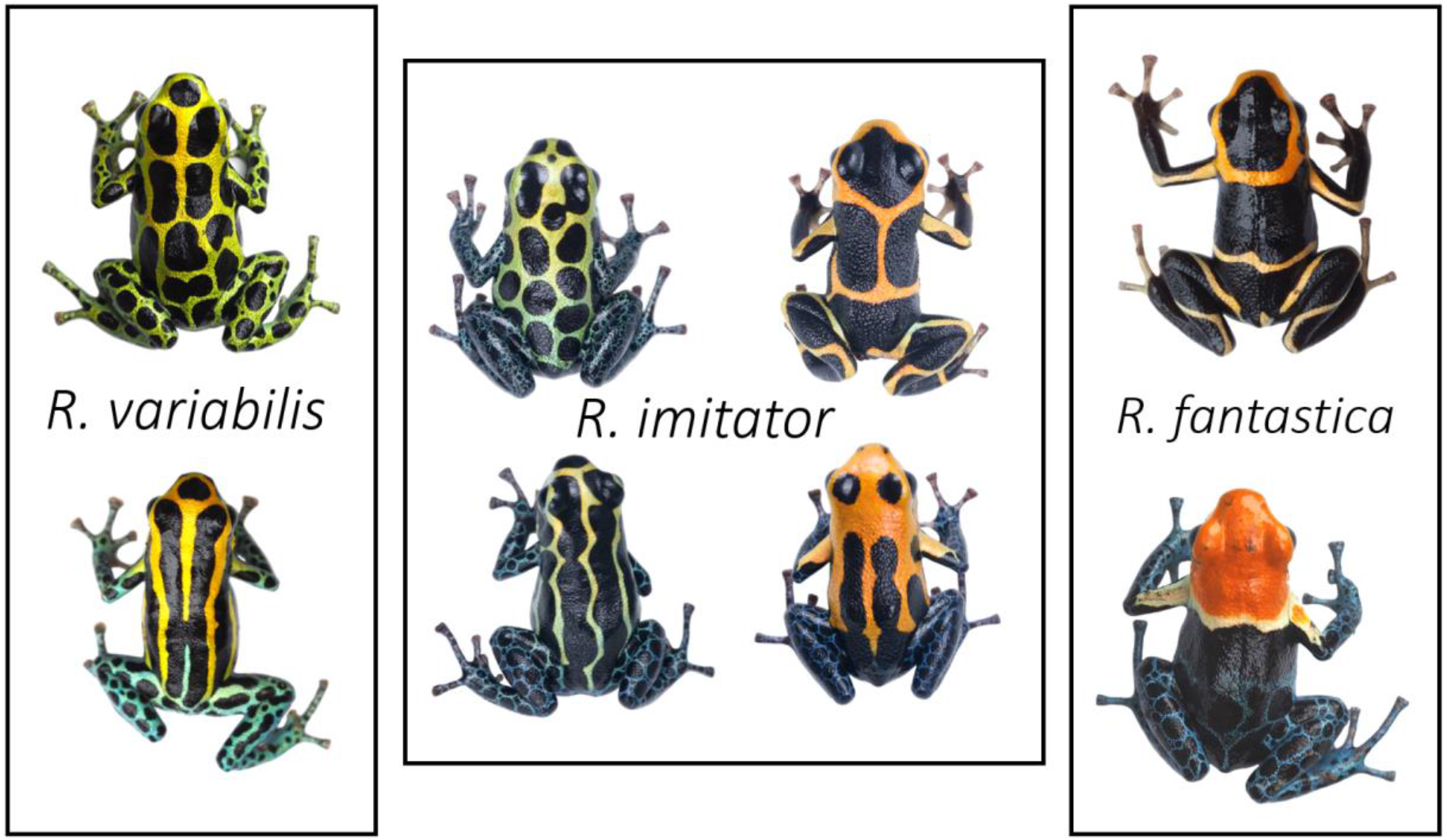
Normative photographs of frogs from the populations used in this study. The center box represents the four mimetic color morphs of *Ranitomeya imitator*. The left exterior box represents the two morphs of *R. variabilis* that *R. imitator* mimics, and the rightmost box represents two color morphs of *R. fantastica* that *R. imitator* has a convergent phenotype with. *R. imitator* photos by AS, *R. fantastica* and *R. variabilis* photos by MC.

**Figure 2.**
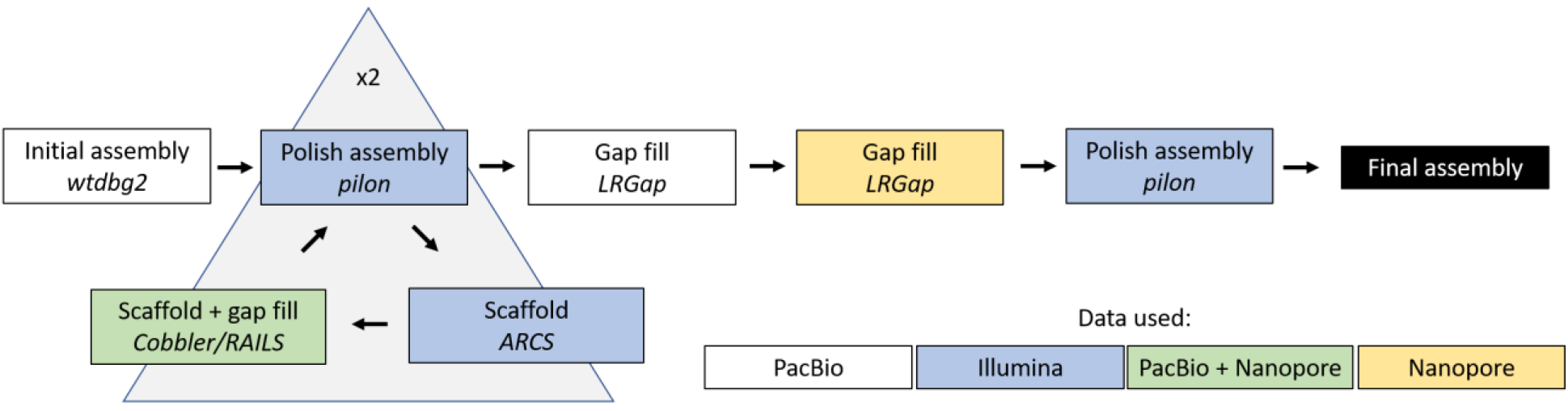
Flowchart of genome assembly approach. Colors of boxes represent the type of data used in that step (see internal legend) and italicized font indicate the program(s) used.

Tadpoles were anesthetized with 20% benzocaine (Orajel), then sacrificed via pithing. Whole skin was removed and stored in RNA later (Ambion) at −20°C until RNA extraction. RNA was extracted from the whole skin using a standardized Trizol protocol, cleaned with DNAse and RNAsin, and purified using a Qiagen RNEasy mini kit. RNA Libraries were prepared using standard poly-A tail purification with Illumina primers, and individually barcoded using a New England Biolabs Ultra Directional kit as per the manufacturer’s protocol. Individually barcoded samples were pooled and sequenced using 50 bp paired end reads on three lanes of the Illumina HiSeq 2500 at the New York Genome Center.

##### *Ranitomeya fantastica* and *Ranitomeya variabilis*

We set up a captive colony consisting of between 6 and 10 wild collected individuals per locality. Three tadpoles per stage (1, 2, 5, 7, and 8 weeks after hatching) were fixed in an RNAlater (Ambion) solution. To do so, tadpoles were first euthanized in a 250 mg/L benzocaine hydrochloride bath, then rinsed with distilled water before the whole tadpole was placed in RNAlater and stored at 4°C for 6h before being frozen at −20°C for long-term storage. Before RNA extraction, tadpoles were removed from RNA later and the skin was dissected off. Whole skin was lysed using a Bead Bug, and RNA was then extracted using a standardized Trizol protocol. RNA libraries were prepared using standard poly-A tail purification, prepared using Illumina primers, and individually dual-barcoded using a New England Biolabs Ultra Directional kit. Individually barcoded samples were pooled and sequenced on four lanes of an Illumina HiSeq X at NovoGene (California, USA). Reads were paired end and 150 base pairs in length.

##### Differential gene expression

We indexed our new *Ranitomeya imitator* genome using STAR version 2.5.4a (Dobin et al., 2013). We removed adaptor sequences from reads using Trimmomatic version 0.39 (Bolger, Lohse, & Usadel, 2014). We then aligned our trimmed reads to our genome using STAR version 2.5.4a (Dobin et al., 2013), allowing 10 base mismatches (--outFilterMismatchNmax 10), a maximum of 20 multiple alignments per read (--outFilterMultimapNmax 20), and discarding reads that mapped at less than 50% of the read length (--outFilterScoreMinOverLread 0.5). We then counted aligned reads using htseq-count version 0.11.3 (Anders, Pyl, & Huber, 2015).

Differential expression analyses were conducted in R version 3.6.0 (Team, 2019) using the package DESeq2 (Love, Anders, & Huber, 2014). Some genes in our annotated genome are represented multiple times, and thus the alignment is nearly to gene level with some exceptions. As a result, when we imported data into R we corrected for this by merging counts from htseq-count into a gene-level count. We filtered out low expression genes by removing any gene with a total experiment-wide expression level ≤ 50 total counts. cDNA libraries for *R. imitator* were sequenced at a different core facility than those of *R. fantastica* and *R. variabilis*, so in order to statistically account for batch effects we analyzed the data from each species independently (combining all species and batch effects in our dataset produces rank deficient models). For each species we created two models using Likelihood Ratio Tests, one which tested the effect of color morph and the other which tested the effect of developmental stage. Both models included sequencing lane, developmental stage, and color morph as fixed effects. We used a Benjamini and Hochberg (Benjamini & Hochberg, 1995) correction for multiple comparisons and used an alpha value of 0.01 for significance. We then extracted data from our models for particular *a priori* color genes that play a role in color or pattern production in other taxa. This *a priori* list was originally used in Stuckert et al. (Stuckert et al., 2019), but was updated by searching for genes that have been implicated in coloration in genomics studies from the last three years. Plots in this manuscript were produced using ggplot2 (Wickham, 2011).

##### Weighted Gene Correlation Network Analysis

To identify networks of genes that interact in response to differences in color morphs, species, or developmental stage, we used weighted gene co-expression network analysis (WGCNA) using the package WGCNA (Langfelder & Horvath, 2008). For the WGCNA analysis, we used the variance stabilizing transformed data produced by DESeq2 at this stage. WGCNA requires filtering out genes with low expression, which we did prior to differential expression analyses. We estimated a soft threshold power (β) that fits our data, by plotting this value against Mean Connectivity to determine the minimum value at which Mean Connectivity asymptotes, which represents scale free topology. For our data, we used β = 10, the recommended minimum module size of 30, and we merged modules with a dissimilarity threshold below 0.25. After module formation, we tested whether the eigenmodules (conceptually equivalent to a first principal component of the modules) were correlated with color morphs, species, or developmental stage at p < 0.05. To examine gene ontology of these modules we then took module membership for each gene within these modules and ranked them in decreasing order of module membership. We then supplied this single ranked list of module membership to the Gene Ontology enRIchment analysis and visuaLizAtion tool (GOrilla), and ran it in fast mode using *Homo sapiens* set as the organism (Eden, Navon, Steinfeld, Lipson, & Yakhini, 2009).

## Results

### Ranitomeya imitator Genome

#### Genome assembly

Our initial contig assembly was 6.77 Gbp in length, had a contig N50 of 198,779 bp, and contained 92.3% of expected tetrapod core genes after polishing with Pilon. After this initial assembly, we conducted two rounds of scaffolding and gap-filling our genome using our data. Our final genome assembly was 6.79 Gbp in length and consisted of 73,158 scaffolds ranging from 1,019-59,494,618 bp in length with an N50 of 397,629 bp (77,639 total contigs with an N50 of 301,327 bp, ranging from 1,019-59,494,618 bp; see Figure 3). A total of 8,149 contigs were placed into scaffolds by our iterative scaffolding and gap-filling, and on average scaffolds contain 1.1 contigs. Based on our BUSCO analysis, the final genome contained 93.0% of expected tetrapod genes. We assembled 69.8% single copy orthologs and 23.2% duplicated orthologs. An additional 1.8% were fragmented and 5.2% were missing (see Supplemental Table 1). The predicted transcriptome from Maker had 79.1% of expected orthologs, 59.4% of which were single copy and complete, 19.7% of which were duplicated, 7.5% of which were fragmented, and 13.4% missing.

**Figure 3.**
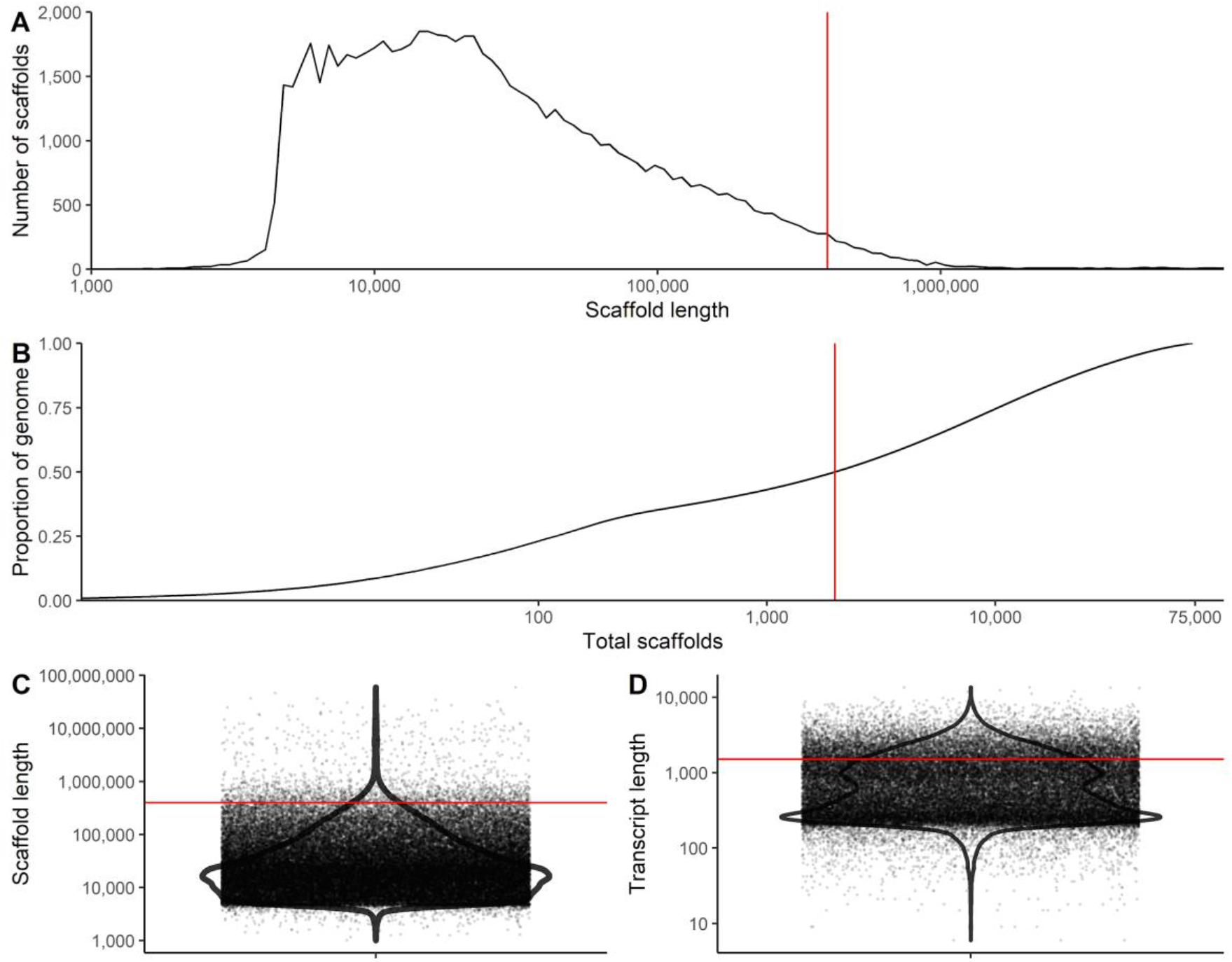
Summary figures of genome assembly. A) Distribution of scaffolds by length. B) The cumulative proportion of the genome represented by scaffolds. Scaffolds were ordered from longest to shortest prior to plotting, therefore in this panel the longest scaffolds are represented in the left portion of the figure. C) Violin plot of the distribution of scaffold lengths in the genome. D) Violin plot of the length of transcripts in the genome as predicted by Maker. Red lines represent N50 in A, C, and D and L50 in B.

#### Repeat elements

Our analyses indicate a high proportion of repeats in the *R. imitator* genome (see Figure 4). Repeat Modeler masked 54.14% of total bases in the genome, of which 50.3% consisted of repeat elements (see Table 1). Repeat masker was unable to classify most of the masked repeat bases (1.7 Gbp representing 25.67% of the genome). Many of the remaining repeats were retroelements (18.36%), nearly 10% of which were LINEs. Just under 8.5% were LTR elements, including 7.45% GYPSY/DIRS1 repeats. Given the quality of repeat databases and the scarcity of amphibian genomic resources in these databases, our results likely represent an underrepresentation of repeats in the genome as a whole. Additionally, many of these repeat classes were long repeats (see Figure 5). For example, the average length of repeats was > 1000 bp in L1 (1093.4 ± 1490.5), L1-Tx1 (1417.3 ± 1372.3), Gypsy (1349.0 ± 1619.3), and Pao (1254.9 ± 1385.8) repeat elements. Summary statistics on the number of instances, range, average length, and standard deviation of range can be found in Supplemental Table 2.

**Figure 4.**
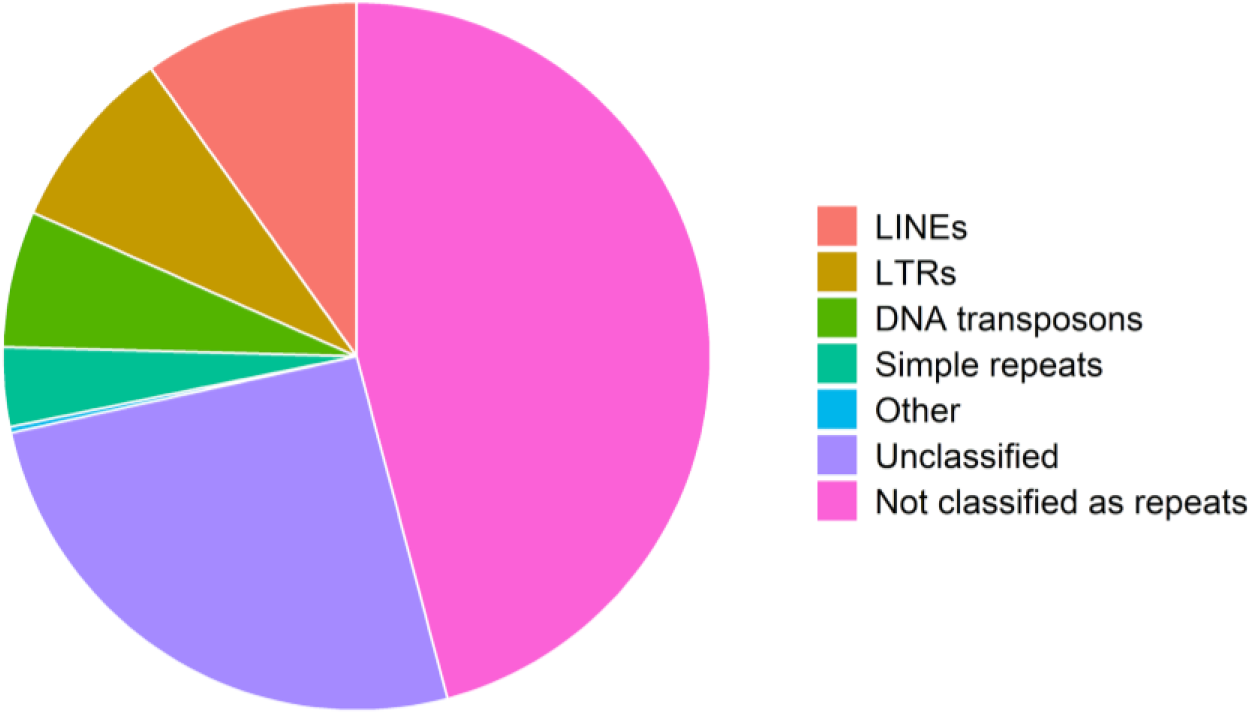
Pie chart of the contents of the *Ranitomeya imitator* genome. Much of the genome is classified as repeat elements, although we were unable to identify many of those.

**Figure 5.**
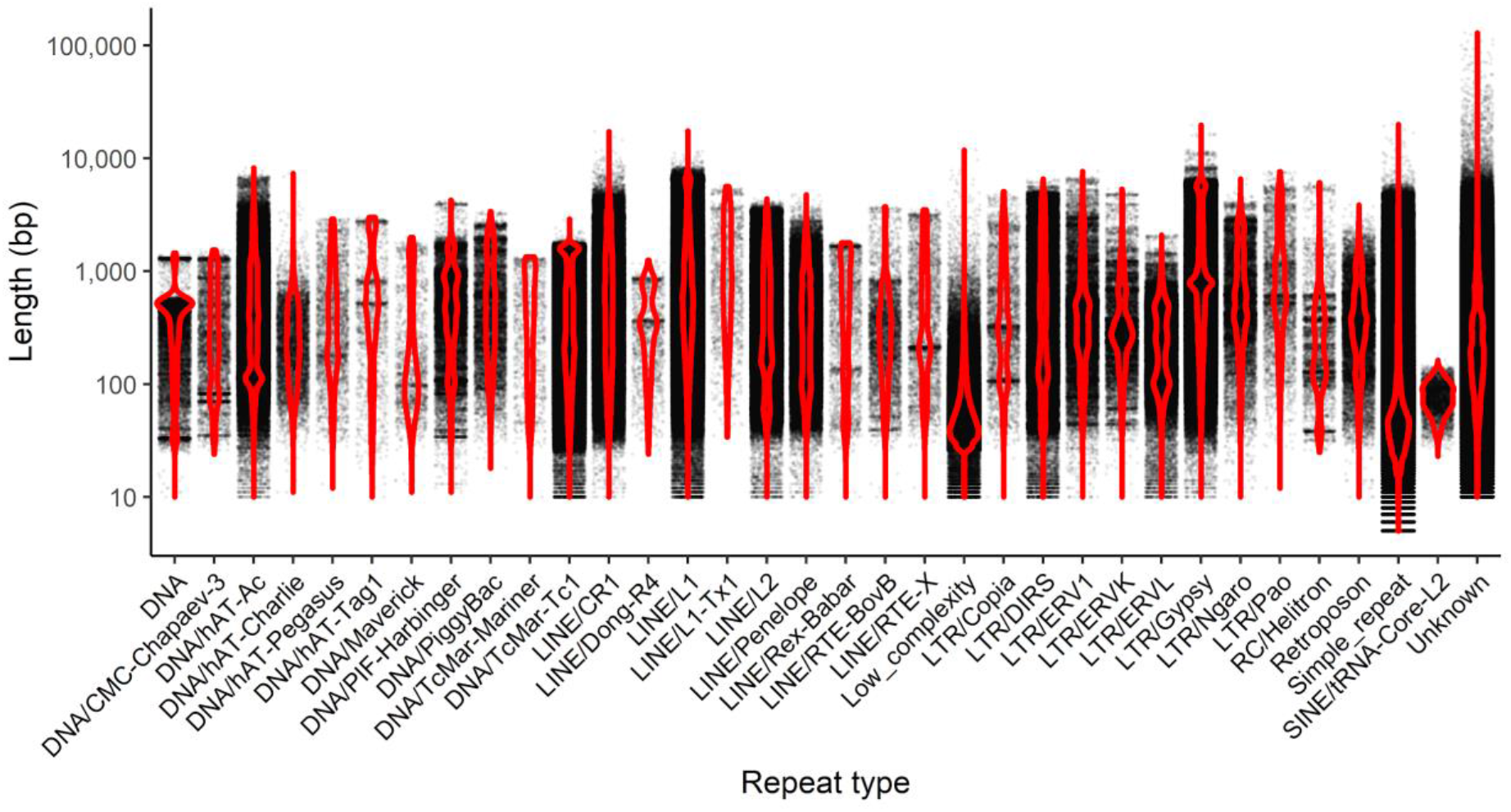
A violin plot of the length of individual repeats in the *R. imitator* genome. Each point represents a single instance of the repeat.

A large proportion of tetrapod orthologs (23.2%) are present in duplicate copies that are spread throughout scaffolds in the genome. These are spread throughout scaffolds in the genome. On average, scaffolds had 1.92 ± 2.93 duplicated orthologs, with a median and mode of 1. When normalized per 10,000 bp, the average was 0.060 ± 0.108 SD duplicated orthologs per 10,000 bp of a scaffold, with a median of 0.032. Intriguingly, the majority of orthologs identified as single copies by BUSCO had average coverage greater than the average genome-wide coverage. Duplicated orthologs, on the other hand, had coverage lower than the average genome-wide coverage. Single copy orthologs had an average coverage of 60.5 ± 29.3 SD, whereas duplicated orthologs had an average coverage of 41.6 ± 31.1 SD, and a two-tailed student t-test revealed a significant difference in coverage between single and duplicate copy orthologs (t_5094_ = 23.965, p-value < 0.0001; See Supplemental Figure 1).

#### Gene expression

We aligned an average of 20.97 million reads (± 6.7 sd) per sample. On average, 73.27% of reads were uniquely mapped (± 4.75% sd). Mapping rates were slightly higher in *R. imitator* than in *R. fantastica* or *R. variabilis*, because the libraries were of slightly higher quality in *R. imitator*. To further test if this was an artifact derived from mapping reads from other species to the *R. imitator* genome, we also mapped these reads to species-specific transcriptome assemblies and found similar results. Thus our mapping rates are driven primarily by the slightly lower quality of the *R. fantastica* and *R. variabilis* cDNA libraries and not by species-specific differences in coding regions. For data on number of reads and mapping rates in each sample please see (Supplemental Table 3).

All gene expression count data can be found in the GitHub repository (https://github.com/AdamStuckert/Ranitomeya_imitator_genome/tree/master/GeneExpression/data). Patterns of gene expression are largely driven by developmental stage (principal component 1; 43% of variation) and species (principal component 2; 13% of variance; see Figure 6). This parallels phylogeny, as *R. fantastica* and *R. variabilis* are more closely related to each other than either are to *R. imitator* (Brown et al., 2011). We then conducted a test for the effect of color morph and developmental stage for each species independently. For a list of all differentially expressed color genes see Supplemental Table 4. For a list of all differentially expressed genes at alpha < 0.01 and alpha < 0.05 see Supplemental Table 5 and 6 respectively.

**Figure 6.**
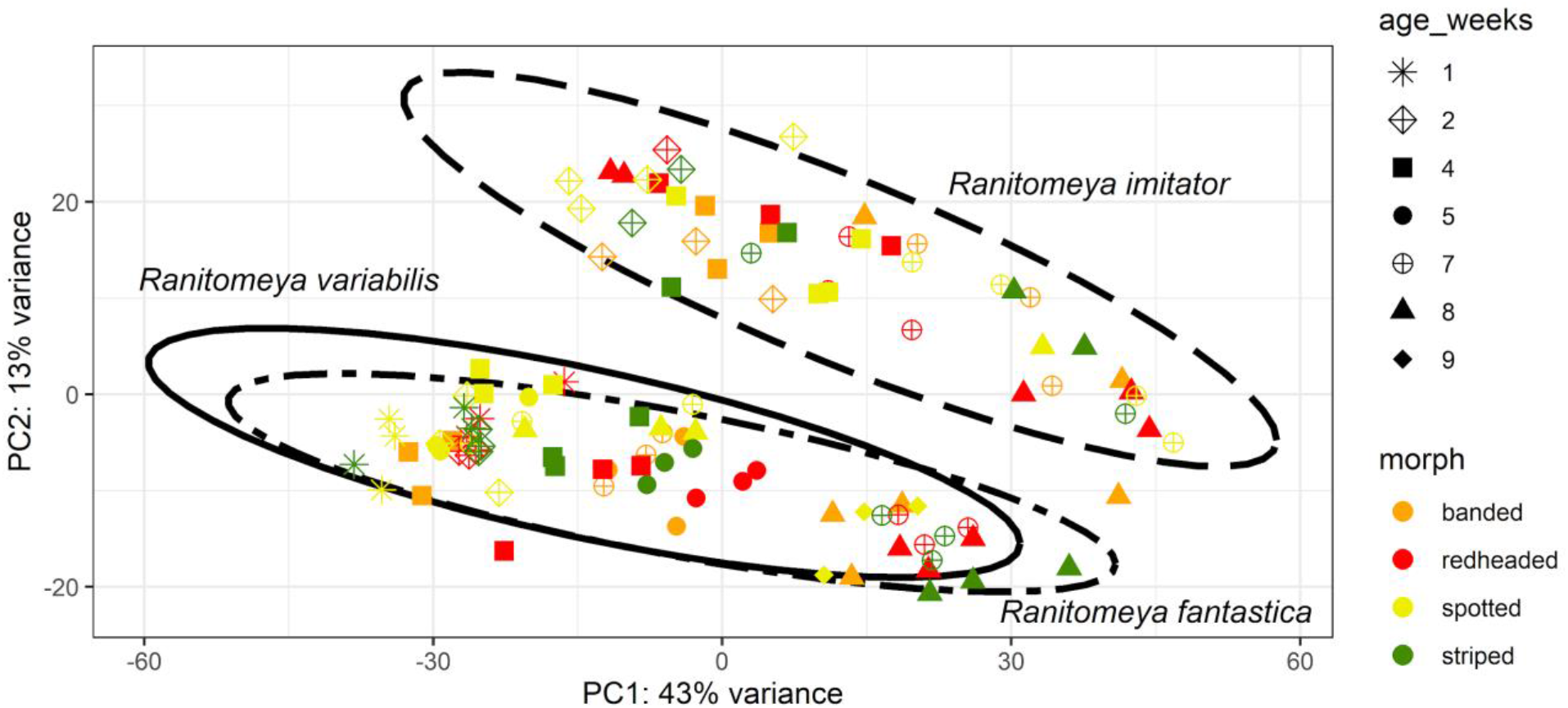
Principal component analysis of gene expression. The axes are labeled with the proportion of the data explained by principal components 1 and 2.

#### Between developmental stage comparisons

In our comparison of developmental stages we found many differentially expressed genes (qvalue < 0.01) in each species (*R. imitator* = 2,039, *R. fantastica* = 2,247, *R. variabilis* = 2,734; Table 2). Most of these are unlikely to be related to color and patterning, although a small fraction of differentially expressed genes (average 3.5%) are related to our *a priori* list of genes that influence the generation of color or patterning in other taxa (*R. imitator* = 92, *R. fantastica* = 106, *R. variabilis* = 77). Amongst genes that were significantly differentially expressed between developmental stages we identified genes related to carotenoid metabolism (e.g., *bco1, retsat, scarb2, ttc39b;* Figure 7), the synthesis of pteridines (e.g., *gchfr, qdpr, pts, xdh;* Figure 8), genes related to melanophore development and melanin synthesis (*dct, kit, lef1, mitf, mlph, mreg, notch1, notch2, sfxn1, sox9, sox10, tyr*, and *tyrp1;* Figure 9), genes putatively related to the production of iridophores and their guanine platelets (e.g., *gart, gas1, paics, pacx2, pax3-a, pnp, rab27a, rab27b, rab7a, rabggta;* Figure 10), and genes related to patterning (*notch1, notch2*).

**Table 2.** Number of differentially expressed genes for each comparison. LRT = likelihood ratio test.

| Species | LRT between morphs |  | LRT between developmental stages |  | Genes differentially expressed between color morphs and developmental stages |
| --- | --- | --- | --- | --- | --- |
|  | Differentially expressed genes | Differentially expressed genes *a priori* color genes | Differentially expressed genes | Differentially expressed genes *a priori* color genes |  |
| *R. imitator* | 1,528 | 39 | 2,039 | 92 | 249 |
| *R. fantastica* | 1,112 | 46 | 2,247 | 106 | 407 |
| *R. variabilis* | 1,065 | 33 | 2,734 | 77 | 574 |

**Figure 7.**
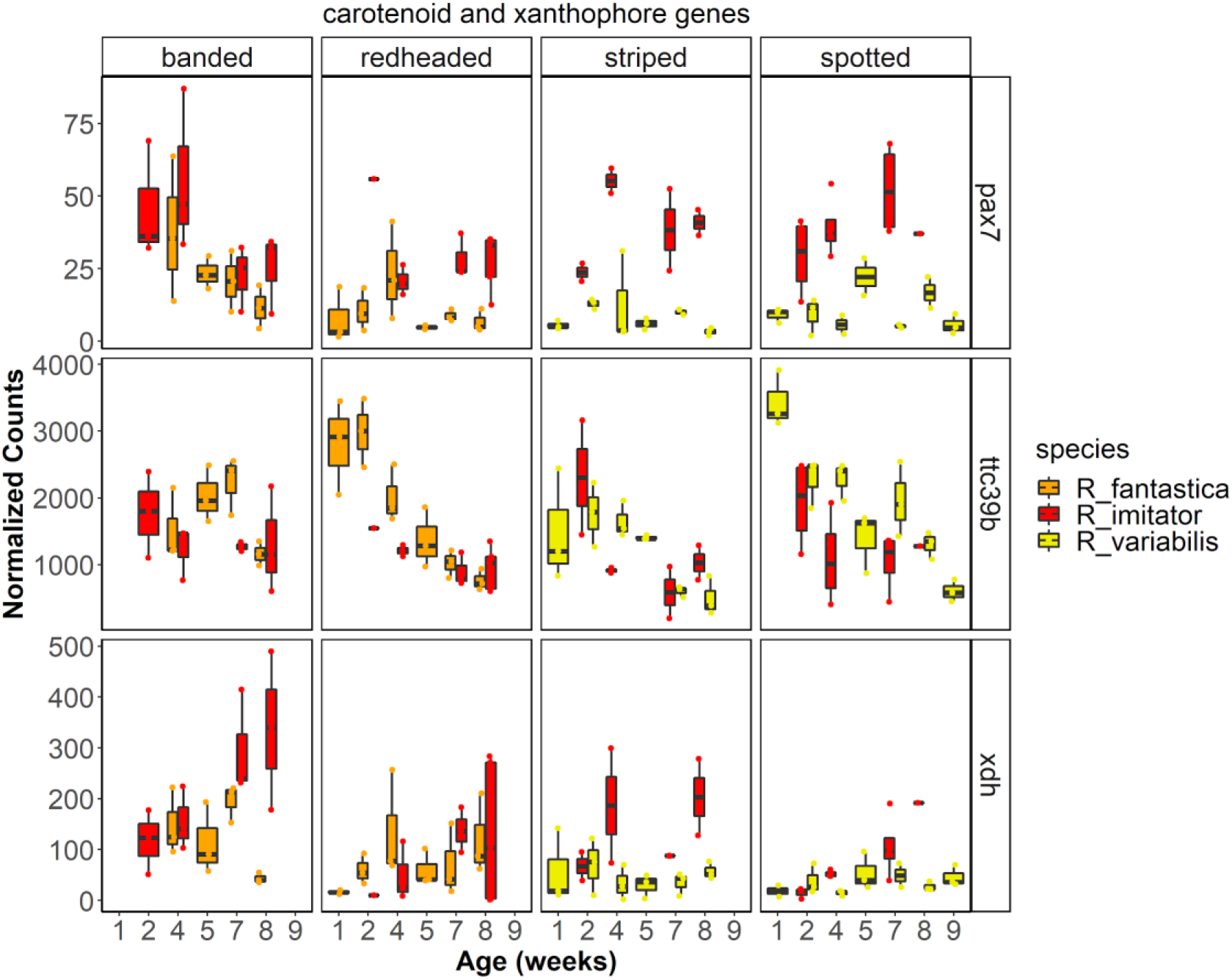
Gene expression for select carotenoid genes.

**Figure 8.**
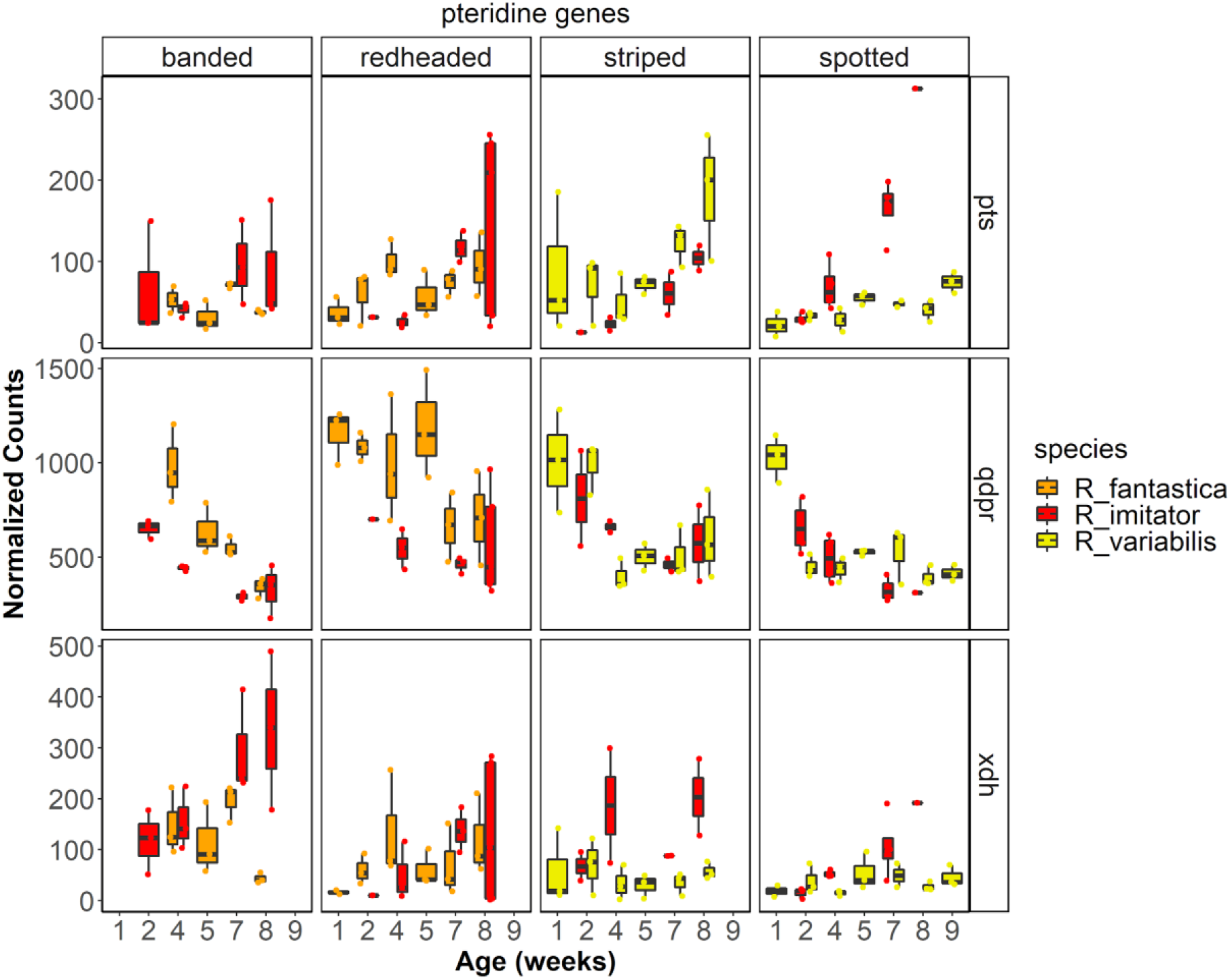
Gene expression for select pteridine genes.

**Figure 9.**
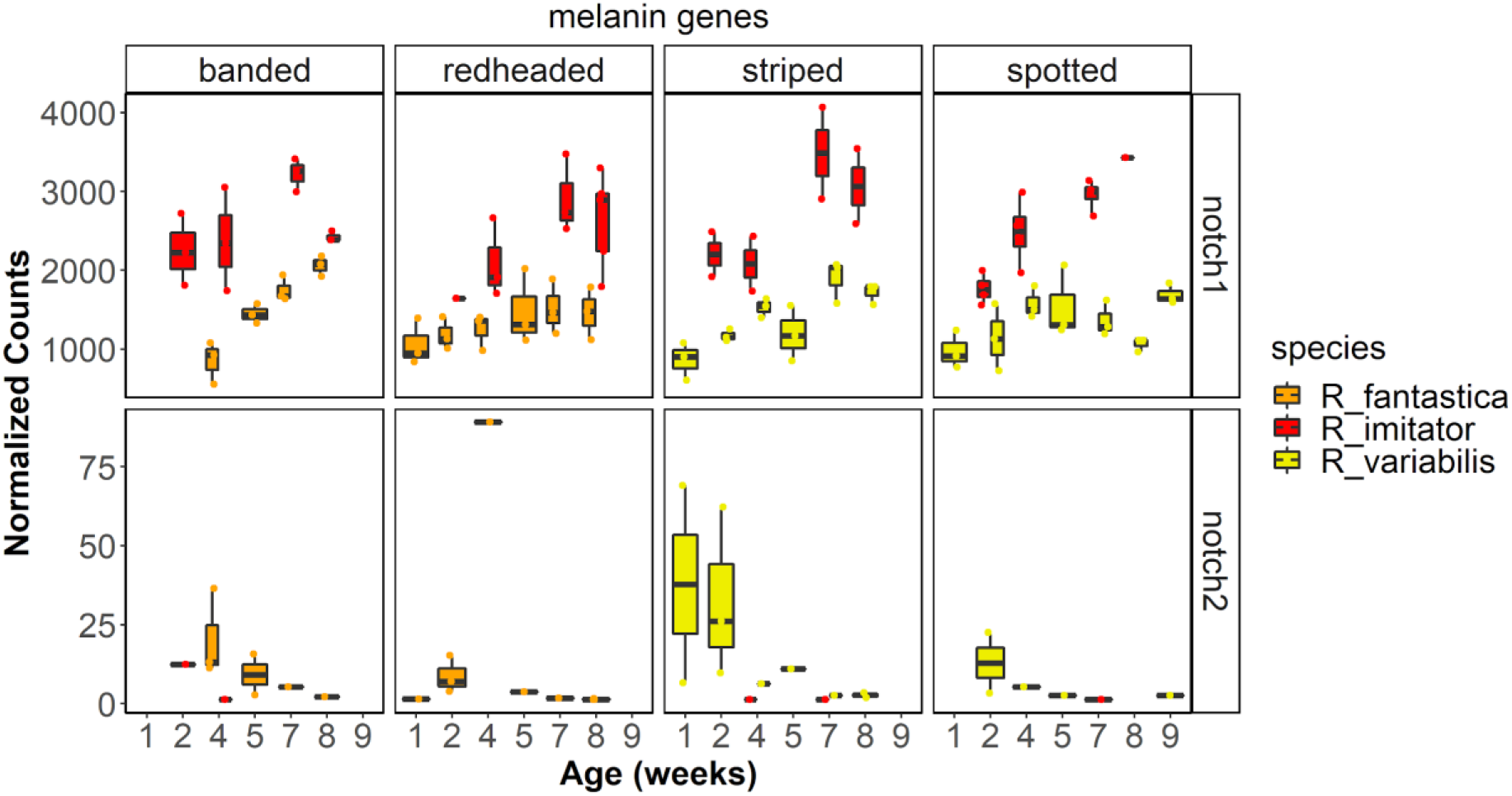
Gene expression for select melanin genes.

**Figure 10.**
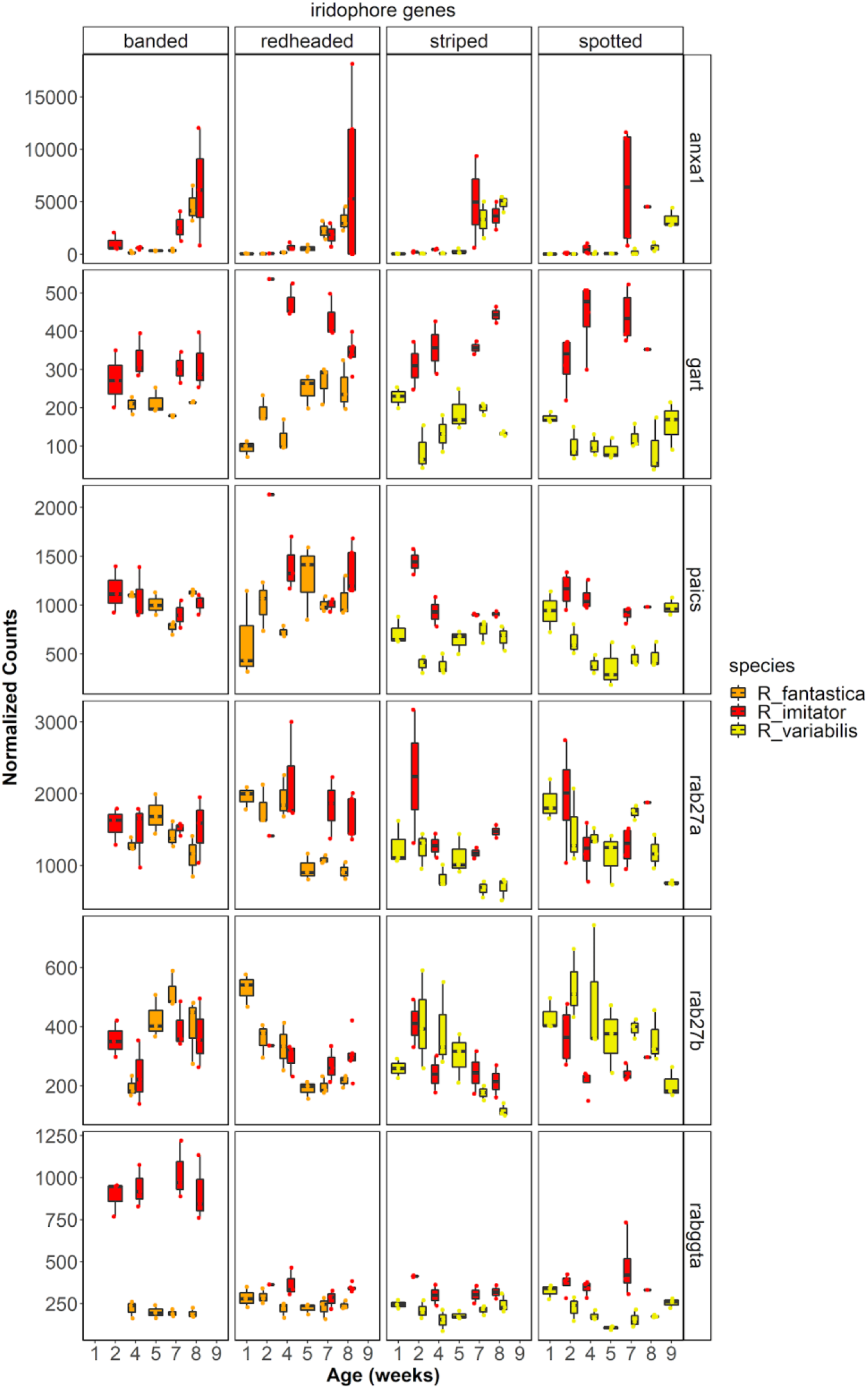
Gene expression for select iridophore genes.

#### Between morph comparisons

In our comparison of color morph we found many significantly differentially expressed genes in each species (*R. imitator* = 1,528, *R. fantastica* = 1,112, *R. variabilis* = 1,065; Table 2). Most of these are unlikely to be related to color and patterning, although a small fraction of differentially expressed genes (average 3.3%) are related to the generation of color or patterning in other taxa (*R. imitator* = 39, *R. fantastica* = 46, *R. variabilis* = 33). Amongst genes that were differentially expressed between color morphs we identified genes related to genes related to carotenoid metabolism or xanthophore production (e.g., *aldh1a1, ttc39b;* Figure 7), the synthesis of pteridines (e.g., *gchfr, qdpr, pts, xdh;* Figure 8), genes related to melanophore development and melanin synthesis (*kit, lef1, mlph, mreg, sfxn1, sox9, sox10;* Figure 9), and genes putatively related to the production of iridophores and their guanine platelets (e.g., *atic, dock7, gart, paics, pacx2, pax3-a, rab27a, rab27b, rab7a, rabggta;* Figure 10).

#### Weighted Gene Correlation Network Analysis

We identified 20 individual gene modules in our expression data using WGCNA (see Figure 11 WGCNA modules). Of these, we identified 14 modules that were significantly correlated with developmental stage; green (r = −0.330; p < 0.0001), midnight blue (r = −0.446; p < 0.0001), purple (r = −0.442; p < 0.0001), greenyellow (r = −0.652; p < 0.0001), light cyan (r = −0.510; p < 0.0001), black (r = −0.268; p = 0.0040), grey 60 (r = −0.405; p < 0.0001), red (r = −0.585; p < 0.0001), light green (r = 0.573; p < 0.0001), blue (r = 0.669; p < 0.0001), cyan (r = 0.734; p < 0.0001), magenta (r = 0.215; p = 0.022), light yellow (r = −0.345; p = 0.00017), and salmon (r = 0.431; p < 0.0001). We identified 10 modules that were significantly correlated with species; green (r = −0.280; p = 0.0025), pink (r = − 0.566; p < 0.0001), purple (r = 0.308; p = 0.000857), greenyellow (r = 0.320; p = 0.000529), light cyan (r = 0.226; p = 0.0156), grey 60 (r = 0.196; p = 0.0366), cyan (r = −0.209; p = 0.0257), magenta (r = 0.304; p = 0.00103), salmon (r = −0.308; p = 0.000869), and grey (r = 0.656; p < 0.0001). Finally, we identified 3 modules that were significantly correlated with color morph, the grey (r = 0.403; p = < 0.0001), pink (r = −0.372; p = < 0.0001), and purple modules (r = 0.190; p = 0.042). Significant GO terms associated with each module that is correlated with species, color morph, and developmental stage are found in Supplemental Tables 7, 8, and 9 respectively.

**Figure 11.**
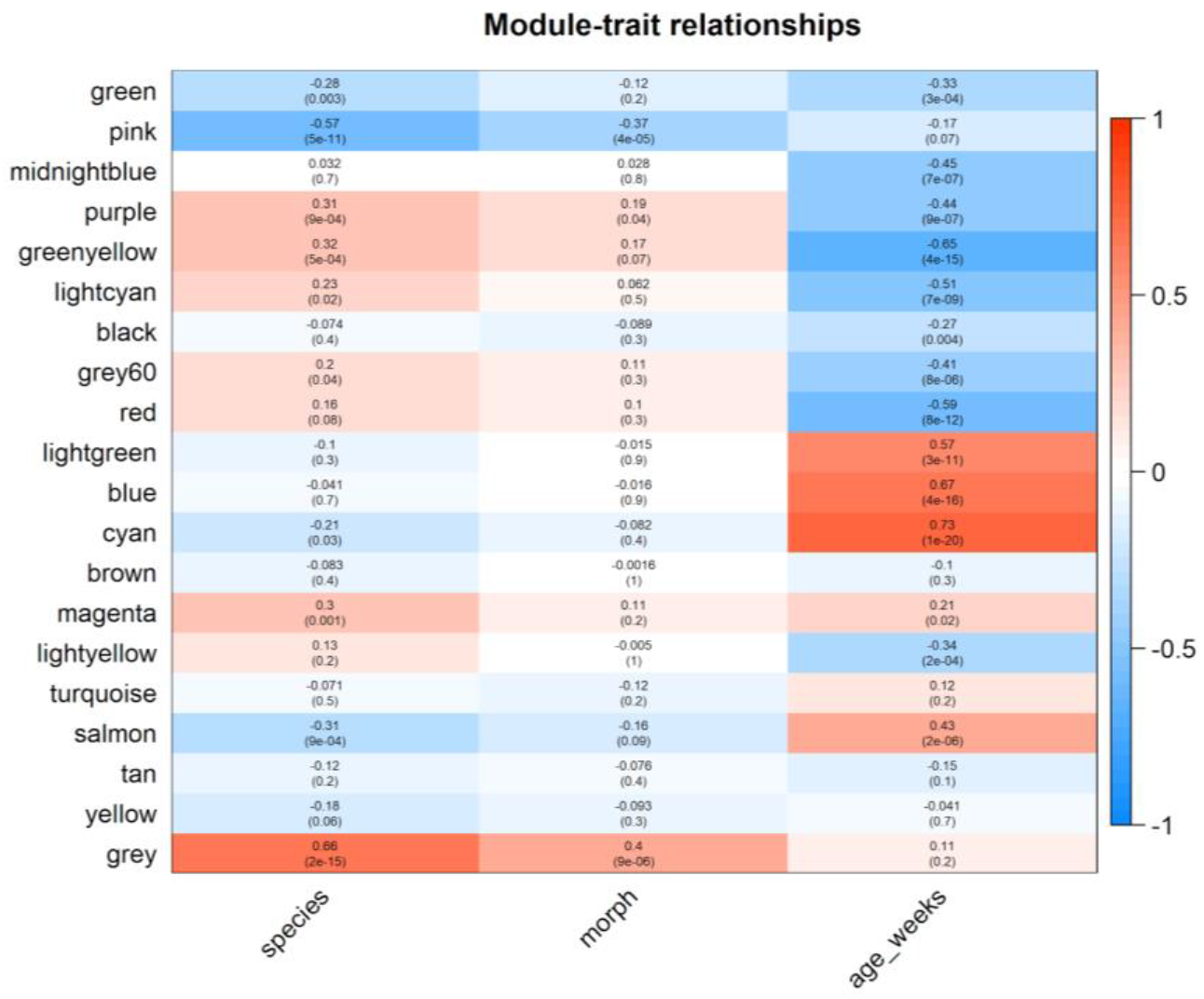
The relationship between WGCNA modules and treatment groups. Within each cell the top number represents the R value (correlation between the grouping and module expression) and the bottom number represents the adjusted p value.

## Discussion

The genetic, biochemical, cellular, physiological and morphological mechanisms that control coloration in mimetic systems are of interest because of their obvious impacts on survival. Despite this, these mechanisms are poorly characterized. Further, genetic mechanisms and genomic resources in amphibians are limited and poorly understood, particularly compared to better known groups like mammals and fish. In this study, we examined how gene expression contributes to differential phenotypes within species in a Müllerian mimicry complex of poison frogs. To do this we assembled a high-quality genome for the mimic poison frog *Ranitomeya imitator*, which we leveraged to conduct gene expression analyses. Here we describe the resulting *R. imitator* genome assembly and highlight key pathways and genes that likely contribute to differential color production within species, illuminating the mechanisms underlying Müllerian mimicry, and providing a rich foundation upon which future research may be built.

### Genome

Our newly assembled *Ranitomeya imitator* genome is a large, high-quality genome with high genic content. This assembled genome is 6.8 Gbp in length and contains 93% of the expected genes according to our BUSCO results. Further, our genome is relatively contiguous with a contig N50 of over 300 Kbp. The *O. pumilio* genome produced with short read technologies had a contig N50 of 385 base pairs and many genic regions were not assembled, presumably because of long intronic regions strewn with repeat elements (Rogers et al. 2018). This dramatic difference in genome contiguity and genic content indicates that long read technologies are, unsurprisingly, critically important and can produce genomes with contiguity spanning large regions, even for species with large genomes containing many long, repetitive regions. Further, the relatively high error rate of current long read technologies does not preclude the ability to assemble and identify genes, as evidenced by our high BUSCO score. This genome is a valuable resource and is well-suited to a variety of future work, especially RNA sequencing analyses like those we present below.

#### *Repeat elements in the* Ranitomeya imitator *genome*

Roughly 50% of our genome assembly was masked as a repeat. This is a large proportion of the genome, but not as large as that of the strawberry poison frog (*O. pumilio*), which was estimated to consist of ~70% repeats (Rogers et al., 2018). Scattered throughout the *R. imitator* genome are a large number of repeat elements which represent over half of the assembled genome. While our assembly of this genome is similar in size to the estimated size of the *Oophaga pumilio* genome, we found a much smaller proportion of particular families of repeat elements (e.g., *Gypsy*) than Rogers et al. (2018) did. Instead, the *R. imitator* genome seems to have a higher evenness of repeat element types spread throughout the genome. A number of the repeat elements that we identified had an average length longer than 1,000 base pairs, including L1 (1,093.4 ± 1,490.5), L1-Tx1 (1,417.3 ± 1,372.3), *Gypsy* (1,349.0 ± 1619.3), and *Pao* (1,254.9 ± 1,385.8). Not only that, but these repeat classes made up a large portion of the genome. Our *R. imitator* genome assembly has less than half the total content represented in the strawberry poison frog genome for many of the most abundant families of repeats in the strawberry poison frog, including *Gypsy* (0.44 Gbp vs 1 Gbp), *Copia* (3 Mbp vs 298 Mbp), *hAT* (97 Mbp vs 255 Mbp), and *Mariner* (0.6 Mbp vs 197 Mbp). However, *R. imitator* has much more total repeat content of *Tc1* (298 Mbp vs 181 Mbp) and a large portion of the *R. imitator* genome consists of unidentified repeats (25%, 1.87 Gbp). Thus, it seems likely that different families of repeats have proliferated between the two poison frog genomes, although the proportion of repeats that we were unable to classify in the *R. imitator* genome prevents us from stating this conclusively.

Our BUSCO analysis also identified a large number of genes that have repeats scattered throughout the genome—23.2% of expected orthologs are present in duplicate copies. These are spread throughout the scaffolds in the genome, and the same repeat region is not found more than at most three times in the genome. The repeats appear almost exclusively in two copies that very rarely appear on the same scaffold. Both copies of a duplicated ortholog only appeared on the same scaffold four times. All of the other duplicated genes were split across scaffolds. Additionally, we found that most single copy orthologs had higher-than-average coverage, whereas most duplicated orthologs had below average coverage. Indeed, relative to single copy orthologs, a larger proportion of duplicated orthologs had particularly low coverage (< 10x).

This evidence, while preliminary, could indicate that some proportion of the large number of duplicated orthologs found in the genome arise from uncollapsed regions of heterozygosity. This is consistent with duplicated orthologs being present in only two copies that often have low coverage. Heterozygosity in a large genome full of long repeats could have led to difficulties in accurately collapsing these regions of the genome, even with long read data. Thus some of the duplicated orthologs were likely incorrectly assembled into multiple copies. Alternatively, this could be driven by a recent duplication event (perhaps chromosomal), which would lead to both multiple copies of orthologs as well as mapping ambiguities due to sequence similarities.

### Gene expression

While the colors and patterns of *Ranitomeya* poison frogs are extremely variable, they always consist of vivid color patches overlaying a background that is largely black in most color morphs. Recent evidence indicates that much of the differences in color in poison frogs are derived from the structure (thickness) and orientation of iridophore platelets (Twomey, Kain, et al., 2020). Additionally, specific pigments that are deposited in the xanthophores, such as pteridines and carotenoids, interact with these structural elements to influence integumental coloration (Duellman & Trueb, 1986; Twomey, Kain, et al., 2020) in the yellow to red ranges of hue. Black and brown coloration is produced by melanophores and the melanin pigments found inside them (Duellman & Trueb, 1986). These data are corroborated by new genomic data that seem to highlight the importance of pigment production and modification genes such as those in the melanin synthesis pathway (Posso-Terranova & Andrés, 2017; Stuckert et al., 2019), pteridine synthesis pathway (Rodríguez et al., 2020; Stuckert et al., 2019) and carotenoid processing pathways (Twomey, Johnson, et al., 2020) for their roles in producing different color morphs in poison frogs.

We identified a large number of differentially expressed genes in our data. This is unsurprising, as we were examining gene expression across a large proportion of tadpole development, a time during which dramatic reorganizations of the body structure are occurring. Many of these genes were differentially expressed throughout development and are likely related to body restructuring rather than coloration *per se.* Nevertheless, we identified a number of very promising candidate color genes that are likely to play a role in the production of mimetic phenotypes in this system.

### Yellow, orange, and red coloration

Yellows, oranges, and reds are determined in large part by the presence of pigments deposited within the xanthophores, the outermost layer of chromatophores in the skin (Duellman and Trueb 1986). These pigments are primarily composed of pteridines and carotenoids, and many studies to date have documented that these pigments play a key role in the production of yellows, oranges, and reds (Croucher, Brewer, Winchell, Oxford, & Gillespie, 2013; Grether, Millie, Bryant, Reznick, & Mayea, 2001; Mcgraw, Nolan, & Crino, 2006; McLean, Lutz, Rankin, Stuart-Fox, & Moussalli, 2017; Obika & Bagnara, 1964). Given the clear importance of xanthophores, pteridines, and carotenoids in the production of yellows, oranges, and reds, we briefly examine the contribution of genes in all three pathways here, beginning with xanthophore production.

#### Xanthophores

While pigments generally determine the color of reds, oranges, and yellows, these pigments all require the structural xanthophores for deposition. We identified a number of key genes that are differentially expressed that are required for the production of xanthophores, notably paired box 7 (*pax7*) and xanthine dehydrogenase (*xdh*). *Pax7* is a transcription factor that is required for establishing and differentiating xanthophores in both embryonic and adult zebrafish (Nord, Dennhag, Muck, & von Hofsten, 2016). *Pax7* was differentially expressed between morphs in *R. fantastica,* notably with higher expression in the orange-banded morph. *Xdh* is sometimes referred to as a xanthophore differentiation marker, as it is found in xanthoblasts and is required for synthesizing the pteridine pigment xanthopterin (Epperlein & Löfberg, 1990; Parichy, Ransom, Paw, Zon, & Johnson, 2000; Reaume, Knecht, & Chovnick, 1991). In our study, this gene exhibited differential expression across development in both *R. imitator* and *R. fantastica*. Additionally, we saw differential expression in this gene between *R. imitator* color morphs, with the highest expression in the orange banded morph. Given their roles in xanthophore differentiation, *pax7* and *xdh* are excellent candidates for production of xanthophores across all *Ranitomeya,* and early differences in expression of these genes could lay the groundwork for markedly different colors and patterns.

#### Pteridine synthesis genes

Genes and biochemical products in the pteridine pathway are important for coloration across a wide variety of taxa, as data from both genetic studies and biochemical assays of pigments point to pteridines as important components of animal coloration. Although a number of genes in the pteridine pathway have been implicated in producing different color patterns, in general the genetic control of pteridine pigmentation is poorly characterized and largely comes from studies of *Drosophila melanogaster (Braasch, Schartl, & Volff, 2007; Kim, Kim, & Yim, 2013).* In this study we found a number of key pteridine synthesis genes that were differentially expressed between color morphs. Prominent amongst these are the aforementioned xanthine dehydrogenase (*xdh*), quinoid dihydropteridine reductase (*qdpr*), and 6-pyruvoyltetrahydropterin synthase (*pts*).

In addition to its role in early xanthophore lineages, *xdh* appears to be highly conserved and its expression plays a role in the production of pterin-based coloration in a variety of taxa such as spiders (Croucher et al., 2013), fish (Parichy et al., 2000; Salis et al., 2019), and the dendrobatid frogs *D. auratus* and *O. pumilio (Rodríguez et al., 2020; Stuckert et al., 2019).* Experimental inhibition of *xdh* causes a reduction in the quantity of pterins, resulting in an atypical black appearance in axolotls (Sally K. Frost, 1978; S. K. Frost & Bagnara, 1979; Thorsteinsdottir & Frost, 1986). Additionally, typically green frogs with deficiencies in the *xdh* gene appear blue due to the lack of pterins in the xanthophores (Sally K. Frost, 1978; S. K. Frost & Bagnara, 1979). As discussed above, *xdh* had the highest expression in the orange banded morph of *R. imitator*. Because *xdh* plays a role in the transformation of pterins into several different yellow pterin pigments such as xanthopterin and isoxanthopterin (Ziegler, 2003), differential expression of *xdh* is a plausible mechanism for the production of orange coloration in the banded *R. imitator*. In sum, *xdh* may function in xanthophore production and/or pterin synthesis and is a likely driver of the production of yellow, orange, and green colors in this mimicry system.

One of the first genes in the pteridine synthesis pathway is 6-pyruvoyltetrahydropterin synthase (*pts*), which is responsible for producing the precursor to both the orange drosopterin pigment as well as yellow sepiapterin pigment. This gene has been implicated in the production of yellow color phenotypes in a variety of systems (Braasch et al., 2007; McLean et al., 2019; Rodríguez et al., 2020), and has been linked to different colors in the poison frog *O. pumilio (Rodríguez et al., 2020)*. While the precise mechanism by which this gene affects coloration is unknown, differential expression of *pts* is often found in relation to yellow or orange skin phenotypes. In our study, we identified significant differential expression throughout development in *R. fantastica* and *R. imitator,* with overall expression of this gene increasing throughout development. Additionally, we see differential expression in *pts* between color morphs of *R. variabilis,* with higher expression in the yellow striped morph, particularly close to the end of development. We did not find differential expression of *pts* in *R. imitator*. This indicates that *pts* may be a critical gene in developing yellow pigmentation in *R. variabilis,* but that it may not play the same role in *R. imitator*. Notably, however, Twomey et al. (Twomey, Johnson, et al., 2020) found that the yellow pterin sepiapterin was only present in very small concentrations in the yellow-striped morph of *R. variabilis*.

Quinoid dihydropteridine reductase (*qdpr*) is another gene involved in the pteridine synthesis pathway and is known to alter patterns of production of the yellow pigment sepiapterin (Ponzone, Spada, Ferraris, Dianzani, & De Sanctis, 2004). We found differential expression in this gene across developmental stages in all three species in this study. Notably, the expression of *qdpr* showed a stark decline over development. *Qdpr* was differentially expressed between color morphs in both *R. imitator* and *R. fantastica,* in both cases with the highest expression in the redheaded morph, although expression was also high in the yellow striped morph of *R. imitator*. The *qdpr* gene was also differentially expressed across populations in another species of poison frog, and was only expressed in light blue or green colored morphs (Stuckert et al., 2019). In combination, this evidence indicates that *qdpr* may be playing a role in the production of pteridine pigmentation in this system and in amphibians generally.

#### Carotenoid genes

Carotenoids are important for both yellow and red coloration across a diversity of life forms (McGraw 2006; Toews et al 2017). While carotenoids are clearly an important class of pigments that broadly influence coloration, few known genes contribute to carotenoid based color differences (e.g., *bco2*, *scarb1*, *retsat*), although this is likely an underestimate of the actual number of genes playing a role in carotenoid synthesis and processing given that we are continuously discovering new genes that play these roles (Emerling, 2018; Twomey, Johnson, et al., 2020). It appears that orange and red colored *Ranitomeya* (outside of *R. imitator)* have slightly higher overall carotenoid concentrations, although these colors are more likely to be derived from the pteridine drosopterin (Twomey et al. 2020). However, different color morphs of *R. imitator* possess similar quantities of carotenoids (Twomey et al. 2020), providing little evidence that carotenoid levels influence coloration in *R. imitator*, but that they may be important in other *Ranitomeya spp*.

We found differential expression of a number of carotenoid genes across development (e.g., *aldh1a1*, *aldh1a2*, *bco1*, *retsat*, *scarb2*). Although we did not find evidence that many known carotenoid genes are a major contributor to differences between specific color morphs in our study, we found differential expression between morphs of some carotenoid genes (e.g., *slc2a11, ttc39b*). We found that a gene recently implicated in carotenoid metabolism, the lipoprotein coding gene tetratricopeptide repeat domain 39B (*ttc39b*), was differentially expressed between color morphs of *R. variabilis* and *R. fantastica* (note the latter finding depends on lowering our stringent alpha value to 0.05). Tetratricopeptide repeat domain 39B is upregulated in orange or red skin in a variety of fish species (Ahi et al., 2020; Salis et al., 2019). Hooper et al (Hooper, Griffith, & Price, 2019) found that this gene is associated with bill color in the long-tailed finch, *Poephila acuticauda*, and they hypothesized that *ttc39b* is being used to transport hydrophobic carotenoids to their deposition site. We found that this gene was most highly expressed in the yellow-green spotted morph of *R. variabilis* and the redheaded morph of *R. fantastica*, consistent with findings from other gene expression studies. Given the repeat occurrence of this gene in pigmentation studies, functional validation of *ttc39b* is likely to be informative.

While we found very few putative carotenoid genes identified in other taxa that were differentially expressed between color morphs we nevertheless identified the oxidoreductase activity gene ontology as the primary gene ontology term in the purple module associated with color morphs. In addition to some of the pteridine genes discussed above, this module contains a number of candidates for carotenoid metabolism, including retinol dehydrogenase and a variety of cytochrome p450 genes. A recent study by Twomey et al. (Twomey, Johnson, et al., 2020) identified *CYP3A80* as a novel candidate for carotenoid ketolase that is preferentially expressed in the livers of red *Ranitomeya sirensis*. Our gene ontology analysis results indicate that there may be additional genes strongly influencing carotenoid synthesis and processing in this system that have not been identified in previous studies.

### Brown and black coloration

Many of the differentially expressed genes in our dataset occurred between developmental stages as tadpoles undergo a complete body reorganization as they prepare to metamorphose. In addition to growth, development, and metamorphosis, during this time tadpoles are producing both the structural chromatophores and the pigments that will be deposited within them. Of these, melanin-based coloration is the best understood aspect of coloration, in no small part because of a long history of genetic analyses in lab mice (Hoekstra, 2006; Hubbard, Uy, Hauber, Hoekstra, & Safran, 2010). As a result, there are a large number of genes that are known to influence the production of melanin, melanophores, and melanocytes. In vertebrates, black coloration is caused by light absorption by melanin in melanophores (Sköld, Aspengren, Cheney, & Wallin, 2016). Melanophores (and the other chromatophores) originate from populations of cells in the neural crest early in development (Park, Kosmadaki, Yaar, & Gilchrest, 2009). The four color morphs of *Ranitomeya* used in this study have pattern elements on top of a generally black dorsum and legs, and therefore melanin-related genes are likely to play a key role in color and pattern, both throughout development and between color morphs.

Given that a large portion of pigmentation develops during development when we sampled individuals, we found that many of our differentially expressed candidate genes are in this pathway. Prominent amongst these genes are *dct, kit, lef1, mc1r, mitf, mlph, mreg, notch1, notch2, sox9, sox10, tyr*, and *tyrp1*, all of which were differentially expressed across development in at least one species. It seems likely these genes are contributing to color production across species given the important roles of each of these genes in melanocyte development and melanin synthesis. In fact, the well-known patterning genes in the notch pathway (e.g., *notch1, notch2*) seem to be important in all three species, as *notch1* was differentially expressed across development in *R. imitator* and *R. variabilis,* and *notch2* in *R. fantastica* (Hamada et al., 2014).

### Blue coloration

Iridophores are largely responsible for blue (and to a lesser extent green) coloration, which is mainly determined by the reflection of light from iridophores (Bagnara et al. 2007). This depends on the presence and orientation of guanine platelets, where thicker platelets tend to reflect longer wavelengths of light (Ziegler 2003; Bagnara et al. 2007; Saenko et al. 2013). Iridophores are a key component of white, blue, and green coloration. Recently, Twomey et al. (2020) found that variation in coloration in *Ranitomeya* and related poison frogs is largely driven by a combination of the orientation and thickness of the guanine platelets in iridophores. Using a combination of electron microscopy, biochemical pigment analyses, and modeling of the interaction between structural elements in the integument and pigmentation within chromatophores they found that much more of the variation in coloration (hue) was driven by differences in the guanine platelet thickness of iridophores than expected. We discuss our findings in light of this recent work, with respect to how they can inform future work in conjunction with the results of Twomey et al. (2020).

The precise mechanisms underlying the development of iridophores and the size and orientation of the guanine platelets are unknown. However, previous work has suggested that ADP Ribosylation Factors (ARFs), which are ras-related GTPases that control transport through membranes and organelle structure and Rab GTPases, are likely important in determining the size and orientation of iridophores (Higdon et al. 2013). We found a number of these genes to be differentially expressed between developmental stages (*arf6, dct, dgat2, dock7, dst, edn3, erbb3, impdh2, paics, psen1, rab27a, rab27b, rabggta*) or color morphs (*anxa1, dock7, dst, erbb3, gart, gne, paics, rab1a, rab27a, rab27b, rab7a, rabggta*) in our study. We also found differential expression of a number of genes that are known to impact guanine or purine synthesis throughout development *(adsl, gart, gas1, fh, qdpr*) and between color morphs (*atic, psat1, qdpr*). A number of these genes *(adsl, dct, dock7, gart, fh, qdpr, psat1, rabggta*) have been implicated in previous work in dendrobatids (Rodríguez et al., 2020; Stuckert et al., 2019) or in other taxa. The genes that have been implicated in previous studies are clearly good candidates for future study. Additionally, the ARFs and Rab GTPases that we identified and are upregulated in iridophores relative to other chromatophore types in fish (e.g., Higdon et al. 2013) are also good candidates. Unfortunately, our understanding of how these genes affect the development of iridophores is limited, and thus more targeted examinations of iridophore production and pigment production are needed. Notably, a number of epidermis-structuring genes (such as those in the krt family) have been implicated in the production of structural colors (e.g., *krt1, krt2*), although more evidence is needed to verify their role in coloration (Burgon et al., 2020; Cui et al., 2016; McGowan, Aradhya, Fuchs, de Angelis, & Barsh, 2006; Stuckert et al., 2019). We identified a number of these that are differentially expressed between color morphs (e.g., *krt1, krt2, krt10, krt14, krt5*). Genes that influence keratin, and organization of the epidermis generally, are good candidates for the production of different colors, as they may produce structural influences on color (via reflectance) in a manner that parallels what we see from guanine platelets. Keratins are known to influence the distribution and arrangement of melanosomes, with an impact on the ultimate color phenotype of animals (Gu & Coulombe, 2007a, 2007b). The combination of differential expression between colors and/or color morphs in multiple expression studies and a plausible mechanism of color modification suggest that genes in the keratin family may well be crucial for producing color differences, although detailed analyses would need to be conducted to confirm this.

#### Conclusion

In this study we examined the molecular mechanisms by which mimetic phenotypes are produced in a Müllerian mimicry system. Through our efforts, we have produced the first high-quality poison frog genome, a 6.8 Gbp, contiguous genome assembly with good genic coverage. We leveraged this to examine gene expression in the skin throughout development of four comimetic morphs from three species of *Ranitomeya*. We identified a large number of genes related to melanophores, melanin production, iridophore production, and guanine synthesis that were differentially expressed throughout development, indicating that many of these are important in the production of pigmentation, although not color morph specific coloration. Genes related to xanthophore production, carotenoid pathways, melanin production and melanophore production were rarely differentially expressed between color morphs, however those genes that were differentially expressed may be critically important in producing polytypic differences within species that drive mimetic phenotypes. Our results indicate that divergence between color morphs seems to be mainly the result of differences in expression and/or timing of expression, but that convergence for the same colour pattern may not be obtained through the same changes in gene expression between species. We identified the importance of the pteridine synthesis pathway in producing these different color morphs across species. Thus, specific production of colors are likely strongly driven by differences in gene expression in genes in the pteridine synthesis pathway, and our data indicate that there may be species-specific differences in this pathway used in producing similar colors and patterns. Further, we highlight the potential importance of genes in the keratin family for producing differential color via structural mechanisms.

## Supporting information

SupplementalTable_1_GenomeAssemblyMetrics

SupplementalTable_2_RepeatSummaryStats

SupplementalTable_3_ReadMappingData

SupplementalTable_4_AllDEColorGenes_alpha0.01

SupplementalTable_5_AllDEGenes_alpha0.01

SupplementalTable_6_AllDEGenes_alpha0.05

SupplementalTable_7_WGCNA_GO_species_results

SupplementalTable_8_WGCNA_GO_morph_results

SupplementalTable_9_WGCNA_GO_developmentalstages_results

## Funding

Funding for this project was provided by an East Carolina University Thomas Harriot College of Arts and Sciences Advancement Council Distinguished Professorship and NSF DEB 165536 to KS, NSF DEB 1655585 to MDM, and XSEDE XRAC MCB110134 to MDM and AMMS.

## Acknowledgements

We are grateful to many individuals for their help with frog husbandry in the lab, including but not limited to M Yoshioka, C Meeks, A Sorokin, K Weinfurther, R Sen, N Davison, M Johnson, M Pahl, N Aramburu, M Tuatanama, R Mori-Pezo, J Richard and S Gallusser. We are also grateful to Laura Bauza-Davila for her work doing RNA extractions, and Andrew Lang for guidance converting RNA to cDNA and preparing samples for sequencing.

## Author contributions

Designed research: AMMS, MC, MM, RN, KS, MDM

Performed research: AMMS, MC, MM, KS, TL

Analyzed data: AMMS, TLP

Contributed to writing: AMMS, MC, MM, TLP, TL, RN, KS, MDM

## Appendix

## SUPPLEMENTAL METHODS

## Genome assembly

## Annotations using Maker

We annotated our genome using Maker version 3.01.02 (Campbell et al., 2014). We used transcript evidence from *Ranitomeya imitator* to aid in assembly (“est2genome=1”). These data include: 1) a developmental series of tadpole skin across color morphs of captive bred *R. imitator* (this paper, see below), 2) liver, skin, and intestine samples from six different wild *R. imitator* populations (Stuckert et al. unpublished data), and 3) brain samples from captive bred *R. imitator* (Geralds et al. unpublished data). The developmental series data were used in order to accurately annotate genes that are expressed in the skin at different time points in order to target color genes. The addition of data from wild frogs and a variety of other tissue types were used to provide additional transcript evidence in an effort to recover more genes after annotation.

## Transcriptome assemblies

## Developmental series

We used data from the imitator developmental series we analyzed in this paper to make transcriptome across developmental time points in the skin. In order to generate an initial reference transcriptome we assembled 40 M randomly subsampled forward and reverse reads sampled across morphs and time points using seqtk (https://github.com/lh3/seqtk) and used the Oyster River Protocol version 1.1.1 (MacManes 2017) to assemble this dataset. Evidence indicates that there is a substantial diminishment of returns in terms of transcriptome assembly completeness from using over 20-30 million reads (MacManes 2017). Initial error correction was done using RCorrector 1.01, followed by adaptor removal and quality trimming using trimmomatic version 0.36 at a Phred score of ≤ 3 (Bolger et al. 2014) since overly aggressive quality trimming has been shown to reduce assembly completeness (MacManes 2014). The Oyster River Protocol (MacManes 2017) assembles a transcriptome by merging multiple assemblies constructed using a series of different transcriptome assemblers and kmer lengths. We constructed the Independent assemblies with Trinity version 2.4.0 (Grabherr et al. 2011), Shannon version 0.0.2 (Kannan et al. 2016), and SPAdes assembler version 3.11 using 35-mers (Bankevich et al. 2012). This deviates slightly from the Oyster River Protocol specified in MacManes (2017), which specifies kmer lengths of 55 and 75 for SPAdes assemblies, but that exceeds our 50 bp sequencing read length. We then merged these individual assemblies using OrthoFuser (MacManes 2017). Finally, we assessed transcriptome quality using BUSCO version 3.0.1 (Simão et al. 2015) and TransRate 1.0.3 (Smith-Unna et al. 2016).

## Transcriptomes from wild frogs of multiple populations

We *de novo* assembled transcriptomes from six populations along a transition zone from the orange banded morph of *R. imitator* (*R. summersi* mimic) through the yellow striped morph (lowland *R. variabilis* mimic). This includes two pure orange banded populations, two pure yellow striped populations, and two admixed populations with highly variable phenotypes. For each population we randomly chose one individual, concatenated Illumina HiSeq 4000 reads from the liver, intestines, and dorsal skin into a single readset per population and then used the Oyster River Protocol version 2.2.8 (MacManes, 2018) to assemble population specific *de novo* transcriptomes. This is similar to the approach above in the section “*Developmental series*”, with some minor differences which we detail here. First, we assembled individual assemblies using Trinity version 2.8.5 (Grabherr et al., 2011), two iterations of SPAdes version 3.13.1 (Bankevich et al., 2012) with kmer values of 55 and 75 respectively, and finally Trans-ABySS version 2.0.1 (Robertson et al., 2010). These individually built transcriptomes were then merged together using OrthoFuser (MacManes 2018). Unique contigs which were dropped in Orthofuser were recovered using a reciprocal blast search of the final assembly against the individual assemblies for unique contigs. We removed all contigs with expression lower than one Transcript Per Million (TPM) using the TPM=1 flag in the ORP. Contigs that were dropped due to low expression but which likely represent expressed genes were recovered by blasting these against the UniProt database. Finally, transcriptome quality was assessed using BUSCO version 3.0.1 (Simão et al. 2015) and TransRate 1.0.3 (Smith-Unna et al. 2016).

## Brain transcriptome assemblies

Geralds et al. (unpublished data) conducted an experiment looking at the effects of parental behaviors and tadpole begging on gene expression in the brains of adult *Ranitomeya imitator*. We randomly chose one individual from individuals that were interacting with begging tadpoles, and those that were not. We concatenated these data into a single forward and reverse read file, then used the Oyster River Protocol version 2.2.8 (MacManes, 2018) to assemble the brain transcriptome. These details are the same as above in the “*Transcriptomes from wild frogs of multiple populations”* section.

## Gene expression

## Ranitomeya imitator

The initial breeding stock of *Ranitomeya imitator* was purchased from Understory Enterprises, LLC (Chatham, Canada). Frogs used in this project represent captive-bred individuals sourced from the following wild populations: Baja Huallaga (yellow-striped morph), Sauce (orange-banded), Tarapoto (green-spotted), and Varadero (redheaded; see Figure 1). Frogs from these populations were collected by Understory and captive bred in Peru prior to shipping captive-bred frogs to a breeding facility in Canada. We purchased individuals that were captive bred in Canada. The frogs used in this study have a similar phenotype to those of individuals found in captivity from their source populations. As with any captive stock sourced from the wild, there is likely an initial bottleneck and corollary reduction in overall heterozygosity, however morphs were not selectively bred to produce desired phenotypes.

Breeding *R. imitator* pairs were placed in 5-gallon terraria containing small (approximately 13 cm) PVC pipes capped on one end and filled halfway with water. We removed tadpoles from the tanks after the male transported them into the pools of water and hand reared them. Although in the wild female *R. imitator* feed their tadpoles unfertilized eggs, tadpoles can survive and thrive on other food items (Brown et al. 2008). We raised experimental tadpoles on a diet of Omega One Marine Flakes fish food mixed with Freeze Dried Argent Cyclop-Eeze, which they received three times a week, with full water changes twice a week until sacrificed for analyses at 2, 4, 7, and 8 weeks of age. At two weeks, tadpoles are limbless, patternless, and a light gray color with two dark black eyeballs. At 4 weeks tadpoles are a slightly darker gray and have back limb buds. Tadpoles had developed their pattern and some coloration as well as reached the onset of metamorphosis at around week 7, and had metamorphosed, were resorbing the tail, and had their froglet patterns at 8 weeks old. Pattern development continues as juveniles and subadults frogs as they grow into the ultimate pattern they possess as adults. Our four sampling periods correspond to roughly Gosner stages 25, 27, 42, and 44 (Gosner 1960). Tadpoles were raised in a homogenous environment, and thus tadpoles were generally at the same developmental stage at the point of sacrifice. Due to inherent variation, some individuals may have been one Gosner stage off the norm. We sequenced a minimum of three individuals at each time point from the Sauce, Tarapoto, and Varadero populations (except for Tarapoto at 8 weeks), and two individuals per time point from the Huallaga population. Individuals within the same time points were sampled from different family groups (Table 1).

Tadpoles were anesthetized with 20% benzocaine (Orajel), then sacrificed via pithing. Whole skin was removed and stored in RNA later (Ambion) at −20°C until RNA extraction. RNA was extracted from the whole skin using a standardized Trizol protocol, cleaned with DNAse and RNAsin, and purified using a Qiagen RNEasy mini kit. Libraries were prepared using standard poly-A tail purification with Illumina primers, and individually barcoded using a New England Biolabs Ultra Directional kit as per the manufacturer’s protocol. Individually barcoded samples were pooled and sequenced using 50 bp paired end reads on three lanes of the Illumina HiSeq 2500 at the New York Genome Center. This yielded on average 24.45M reads per library ± 8.6M sd (range: 10.1-64.M).

## *Ranitomeya fantastica* and *Ranitomeya variabilis*

We set up a captive colony consisting of between 6 and 10 wild collected individuals per locality, which was maintained at the Tarapoto Research Center (INIBICO. jr. Ventanilla s/n Sector Ventanilla. Banda de Shilcayo. San Martin, Peru). Male-female pairs were placed into individual terrariums which were misted with rainwater and fed with fruit flies daily. Artificial egg deposition sites consisting of short sections of PVC pipe (~10 cm in length) were positioned within each terrarium and we checked terraria for eggs biweekly. When eggs were found, they were transferred into petri dishes to monitor their development. Upon hatching, tadpoles were placed individually into 25 ml plastic cups filled with rain water and fed daily with a pinch of a 50/50 mix of spirulina and nettle leaf powder. Three tadpoles per stage (7, 14, 35, 49 and 56 days after hatching) were fixed in an RNAlater (Ambion) solution. To do so, tadpoles were first euthanized in a 250 mg/L benzocaine hydrochloride bath, then rinsed with distilled water before the whole tadpole was placed in RNAlater in a 2.5ml Eppendorf tube. The Eppendorf tube with sample in RNAlater was then stored at 4°C for 6h before being frozen at −20°C for long-term storage. This protocol was approved by the Peruvian Servicio Forestal y de Fauna Silvestre through the authorization number 232-2016-SERFOR/DGGSPFFS.

Tadpoles were stored whole in RNAlater at −20°C until RNA extraction. Tadpoles were removed from RNA later and the skin was dissected off. Whole skin was lysed using a Bead Bug, and RNA was then extracted using a standardized Trizol protocol. Libraries were prepared using standard poly-A tail purification, prepared using Illumina primers, and individually dual-barcoded using a New England Biolabs Ultra Directional kit. Individually barcoded samples were pooled and sequenced on four lanes of an Illumina HiSeq X at NovoGene (California, USA). Reads were paired end and 150 base pairs in length.

## SUPPLEMENTAL RESULTS

## Genome sequence data

We produced a large set of sequence data (Table XX). This includes approximately 228,115,533,900 10X bases, 22,076,177,973 Oxford Nanopore bases, and 297,076,730,441 PacBio bases.

**Appendix Table 1.** Sample sizes for gene expression of model species *R. fantastica* and *R. variabilis*.

| Species | Morph | Age (weeks) | n |
| --- | --- | --- | --- |
| *Ranitomeya fantastica* | Redheaded | 1 | 3 |
|  |  | 2 | 3 |
|  |  | 4 | 3 |
|  |  | 5 | 3 |
|  |  | 7 | 3 |
|  |  | 8 | 3 |
|  | White-Banded | 4 | 3 |
|  |  | 5 | 3 |
|  |  | 7 | 3 |
|  |  | 8 | 3 |
| *Ranitomeya variabilis* | Spotted | 1 | 3 |
|  |  | 2 | 3 |
|  |  | 4 | 3 |
|  |  | 5 | 3 |
|  |  | 7 | 3 |
|  |  | 8 | 3 |
|  |  | 9 | 3 |
|  | Striped | 1 | 3 |
|  |  | 2 | 3 |
|  |  | 4 | 3 |
|  |  | 5 | 3 |
|  |  | 7 | 3 |
|  |  | 8 | 3 |

**Supplemental Figure 1.**
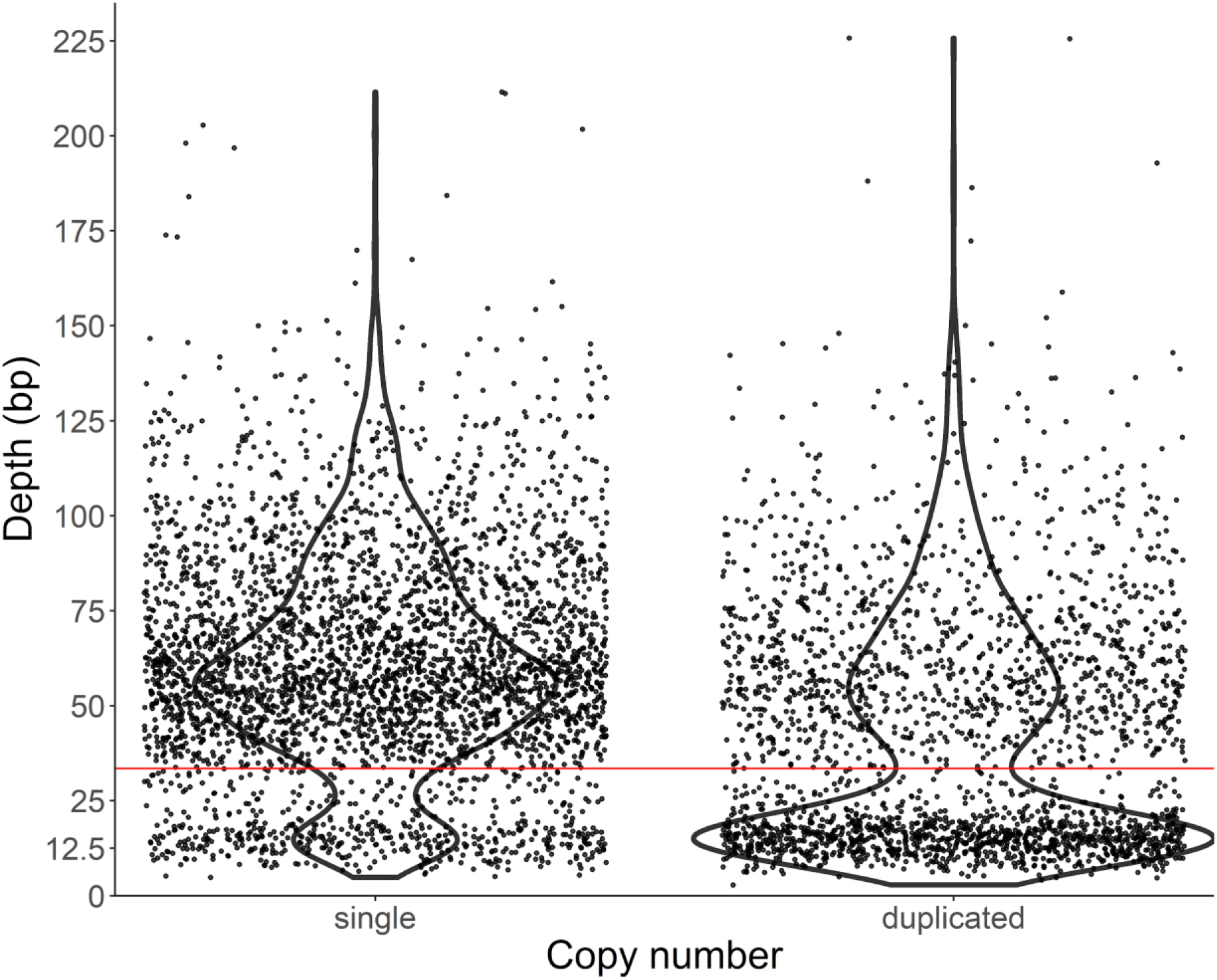
Average coverage for each ortholog in the tetrapoda database. Orthologs are split by whether they are identified by BUSCO as present in the assembly as a single copy or duplicated copies.

## Notes

https://www.ebi.ac.uk/ena/browser/view/PRJEB28312

### Competing Interest Statement

The authors have declared no competing interest.

### Summary of Updates

A previous version of this manuscript examined gene expression of the skin of four color morphs of the poison frog *Ranitomeya imitator* throughout larval development. This used a *de novo* transcriptome. In this revision, we have: 1) Added a whole genome assembly for *Ranitomeya imitator* and relevant analyses of the assembly. 2) Added gene expression data for two species that *Ranitomeya imitator* mimics: *R. fantastica* and *R. variabilis*. 3) Conducted differential expression analyses for all species to our new *de novo* genome assembly for *Ranitomeya imitator*.

