## SupplementalTable_1_GenomeAssemblyMetrics for "The genomics of mimicry: gene expression throughout development provides insights into convergent and divergent phenotypes in a Müllerian mimicry system"

| Assembly | Genome size (GB) | Contig N50 | Scaffold N50 | %Ns | BUSCO | Change |
| --- | --- | --- | --- | --- | --- | --- |
| Imi.ctg.wtdbg | 6.77 | 198,779 | NA | NA | C:89.8%[S:86.5%,D:3.3%]  F:5.3%,M:4.9% | Initial contig assembly |
| Imitator.1.0 | 6.77 | 198,779 | NA | NA | C:92.3%[S:75.4%,D:16.9%]  F:4.6%,M:3.1% | Polished |
| Imitator.1.1 | 6.78 | 198,779 | 330,635 | 0.02 | C:92.3%[S:74.9%,D:17.4%]  F:4.4%,M:3.3% | 10X scaffolded |
| Imitator.1.2 | 6.79 | 204,382 | 339,129 | 0.01 | C:92.5%[S:75.1%,D:17.4%]  F:4.4%,M:3.1% | Cobbler + RAILS |
| Imitator.1.2.1 | 6.79 | 211,576 | 339,195 | 0.01 | C:92.7%[S:74.5%,D:18.2%]  F:4.3%,M:3.0% | Polished |
| Imitator.1.2.2 | 6.79 | 247,642 | 339,195 | 0.01 | C:92.6%[S:74.5%,D:18.1%]  F:4.3%,M:3.1% | LRgapfilled, ONT data |
| Imitator.1.2.3 | 6.79 | 272,070 | 339,195 | 0.00 | C:92.7%[S:74.5%,D:18.2%]  F:4.3%,M:3.0% | LRgapfilled, PacBio data |
| Imitator.1.3 | 6.79 | 275,328 | 339,195 | 0.00 | C:92.7%[S:74.0%,D:18.7%]  F:4.3%,M:3.0% | Polished |
| Imitator.1.3.1 | 6.79 | 275,328 | 397,353 | 0.01 | C:92.6%[S:73.6%,D:19.0%]  F:4.3%,M:3.1% | 10X scaffolded |
| Imitator.1.3.2 | 6.79 | 275,704 | 397,634 | 0.01 | C:92.6%[S:73.6%,D:19.0%]  F:4.3%,M:3.1% | Cobbler + RAILS |
| Imitator.1.3.3 | 6.79 | 275,704 | 397,634 | 0.01 | C:92.6%[S:73.6%,D:19.0%]  F:4.3%,M:3.1% | Polished |
| Imitator.1.3.4 | 6.79 | 292,624 | 397,633 | 0.01 | C:92.7%[S:73.6%,D:19.1%]  F:4.3%,M:3.0% | LRgapfilled, ONT data |
| Imitator.1.3.5 | 6.79 | 300,673 | 397,633 | 0.01 | C:92.7%[S:73.6%,D:19.1%]  F:4.3%,M:3.0% | LRgapfilled, PacBio data |
| Imitator.1.3.6 | 6.79 | 301,327 | 397,629 | 0.01 | C:92.7%[S:73.6%,D:19.1%]  F:4.3%,M:3.0% | Polished |

Table XX. Genome quality metrics. BUSCO was run against the tetrapoda database of 3,950 genes in genome mode. BUSCO numbers are S: single copy genes, D: duplicated genes, F: fragmented genes, M: missing genes. The change column represents the most recent modification to the genome assembly. Chronological order is from top to bottom.
